# Dimerisation of APOBEC1 is dispensable for its RNA/DNA editing activity and modulates its availability

**DOI:** 10.1101/410803

**Authors:** Martina Chieca, Marco Montini, Francesco Severi, Gaia Lembo, Francesco Donati, Riccardo Pecori, Silvestro G. Conticello

## Abstract

The AID/APOBECs are DNA/RNA deaminases whose mutagenic activity has been linked to cancer. Among them, APOBEC1 physiologically partakes into a complex that edits a CAA codon into UAA Stop codon in the transcript of Apolipoprotein B (APOB), a protein crucial in the transport of lipids in the blood. Catalytically inactive mutants of APOBEC1 have a dominant negative effect on its activity, as they compete for the targeting of the APOB mRNA. Here we titrate APOBEC1-mediated editing in presence of catalytically inactive chimeras and mutants of APOBEC1, and we show that APOBEC1 inability to dimerise is the main determinant for its activity. This property is especially evident in an APOBEC1 mutant (L173A G227A) with increased activity on RNA despite decreased self-interaction. Moreover, dimerisation protects APOBEC1 from degradation and regulates its availability. Considering APOBEC1 capability to target DNA, we demonstrate that increased availability of the protein due to dimerisation leads to increase in the DNA damage induced by APOBEC1. These findings demonstrate that dimerisation, a property common to other APOBECs targeting DNA, might represent another layer in the regulation of this editing enzyme.

**BULLET POINTS:**

- APOBEC1 inability to dimerise is the main determinant for its activity.
- Dimerisation protects APOBEC1 from degradation and regulates its availability.
- Alterations in the balance between monomeric and dimeric APOBEC1 increase DNA damage.

## INTRODUCTION

The AID/APOBECs are deaminases that target endogenous and exogenous genetic material, mainly in the context of the immune processes (1). Their aberrant mutagenic activity has been linked to the onset and progression of genetic alterations in cancer (2). Among them, Apolipoprotein B Editing Complex, catalytic subunit 1 (APOBEC1) targets both DNA and RNA. Overexpression of APOBEC1 in model animals induces cancer (3) and its deficiency in cancer-prone mice leads to a decrease of cancer formation (4). APOBEC1 oncogenic potential is likely dependent on its mutagenic activity (5–7). On the other hand, APOBEC1 RNA targeting has been physiologically associated with the editing of cytosine 6666 in a CAA codon (Q2180) of the Apolipoprotein B (APOB) transcript to generate a Stop codon (8–10). APOBEC1-mediated RNA editing leads to the synthesis of a truncated APOB (APOB48), whose balance with the long isoform (APOB100) in liver and small intestine regulates the formation of lipoprotein particles and the transport of cholesterol in the bloodstream. In the years, a number of other targets have been identified (11–17), suggesting that APOBEC1-mediated RNA editing could be a widespread phenomenon to increase transcriptome heterogeneity: C>U editing can recode transcripts, create novel miRNA targets or disrupt existing ones. While the full-complexity of physiological APOBEC1-mediated RNA editing is still not clear, it is possible that aberrant RNA editing by APOBEC1 could be linked to the onset of disease (16, 18).

Higher-molecular complexes are formed by APOBEC1 in the cell (19–21), likely due to nonspecific interactions. Yet, the core editing complex is defined by APOBEC1 partaking into interactions with either APOBEC1 Complementation Factor (A1CF) or RNA-Binding-Motif-protein-47 (RBM47) (22–25), RNA-binding proteins that target APOBEC1 to the specific mRNAs. APOBEC1-mediated RNA editing is a predominantly nuclear process that involves APOBEC1 shuttling from the cytoplasm to the nucleus (26, 27) and acting as a carrier for A1CF (28).

APOBEC1 is able to form homodimers (29), and this characteristic has been hypothesised to be important for its function. Indeed, APOBEC1 deletion mutants lacking the C-terminal domain are unable to edit RNA and they do not self-interact (30). Contradictory results using constructs with partially deleted C-terminal domains to edit RNA *in vitro* indicate that the relation between RNA editing and dimerisation is not clear (31, 32).

It has been shown that, while a catalytically inactive APOBEC1 mutant unable to coordinate Zinc can act as a dominant negative, mutations or deletions in the C-terminal region do not (30). This suggests that interaction through the C-terminal region with at least one other factor, be it the target RNA or another protein, is necessary for APOBEC1 function.

Here we exploit chimeric and mutant APOBEC1 constructs to analyse the dynamics of APOBEC1-mediated RNA editing in live cells. By testing their interplay in live cells using FACS-based RNA editing assays (33), we demonstrate that dimerisation is dispensable for APOBEC1 activity on RNA and it modulates its availability both for RNA and DNA editing.

## MATERIAL AND METHODS

### Plasmids

The EGFP-APOBEC1 and FLAG-APOBEC1 expression plasmids were described in Conticello et al (34). APOBEC1-EGFP, APOBEC1-EGFP(Y66H), EGFP(Y66H)-APOBEC1, pEGFP mCherry-APOB-EGFP and the A1CF expression plasmids are described in Severi and Conticello (33). The FLAG-tagged A1CF expression plasmid was generated by inserting a FLAG expressing adaptor (CTAGACCATGGACTACAAGGACGATGATGACAAGCTTAAG and TCGACTTAAGCTTGTCATCATCGTCCTTGTAGTCCATGGT) 5’ to the A1CF coding sequence (NheI/SalI). FLAG-tagged RBM47 expression plasmid was generated from PCR-amplifed RBM47 (aaactcgagATGACCGCAGAGGATTCC and aaaggtaccTCAGTATGTCTGGTAGAC primers) inserted using XhoI/KpnI restriction sites in a pEGFP-C3 with a FLAG epitope sequence plasmid replacing the EGFP coding sequence. The untagged rat APOBEC1 construct was generated by excising the coding sequence from the plasmid described in Severi and Conticello (33) (NheI/BglII) and inserting it into a pEGFP-C3 backbone in which the EGFP had been removed (NheI/BamHI). The expression constructs for the catalytic inactive APOBEC1 (E63A) were generated by PCR amplification of the coding sequence in the E63A plasmid described in Severi and Conticello (33) (aaagctagcatgagtTCCGAGACAGGCCCTGTA and aaatgtacaagatcTCATTTCAACCCTGTGGC) and cloned into either pEGFP-C3 or pEGFP-N1 backbones containing a non-fluorescent EGFP mutant (Y66H - to avoid interference with the RNA-editing assay), or a pEGFP-C3 without EGFP.

The mutations to generate the non-dimerising rat APOBEC1 mutants (L173A, G227A, L173A/G227A - with or without a FLAG epitope) were inserted by site directed mutagenesis of the corresponding plasmid (L173A mutation primers: TGGCCAAGGTACCCCCATGCGTGGGTGAGG and CCTCACCCACGCATGGGGGTACCTTGGCCA; G227A mutation primers: CCTGTGGGCCACAGCGTTGAAATGAGATCC and GGATCTCATTTCAACGCTGTGGCCCACAGG). The reporter plasmid for the FACS-based RNA editing assay is reported in Severi and Conticello ((33), Addgene # 112859).

### Cell cultures, transfections, and flow cytometry

HEK293T cells were maintained in DMEM supplemented with 10% FBS, 2mM L-Glutamine and 1mM penicillin/streptomycin at 37°C in 5% CO2.

Transient transfections were carried out using either Lipofectamine LTX (Invitrogen - catalog #15338100) according to the manufacturer’s instructions or PolyEthylenimine (PEI, Polyscience - catalog #23966) following Durocher et al (35) using a DNA:PEI ratio of 1:3. The amount of plasmid constructs used in the titration experiments was determined based on the best separation between the titration induced by the empty plasmid or by the E63A mutant (Supplementary Fig. 1). Proteasomal degradation was assessed by MG132 treatment (10μM) for 16 hours.

FACS analysis was performed on a CytoFLEX S (Beckman Coulter) flow-cytometer after 30-48 hours from transfection.

To assay DNA damage induction by FACS, transfected cells were fixed at 48 hours with 4% formaldehyde solution, permeabilized, and stained using anti-phospho-Histone H2AX (Ser139) (1:1500, Merck Millipore - catalog # 05-636) and Rhodamine Red™-X goat anti-mouse IgG (1:2000 Invitrogen - catalog # R-6393). Nuclei were stained with DAPI. Cells were scraped and resuspended in PBS and then visualised on CytoFLEX S (Beckman Coulter) flow-cytometer.

### Immunofluorescence and microscopy analysis

Transfected cells were plated 24 hours after transfection on coverslips pre-treated with poly-D-Lysine hydroxide and fixed at 48 hours. Following fixation with ice cold methanol, cells were stained using either rabbit anti-A1CF (1:400, Sigma - catalog # HPA037779) or rabbit anti-FLAG (1:800, Cell Signalling – catalog # 2368), together with anti-rabbit IgG conjugated with Alexa Fluor 647 (1:800, Life technologies - catalog # A21246); goat anti-APOBEC1 (1:400, Santa Cruz - catalog # 11739) and donkey anti goat IgG H&L conjugated with Alexa Fluor 647 (1:800, Abcam - catalog # 150131); mouse anti-phospho-Histone H2AX (Ser139) (1:800, Merck Millipore - catalog # 05-636) and Rhodamine Red™-X goat anti-mouse IgG (1:1000 Invitrogen - catalog # R-6393). Nuclei were stained with DAPI.

Cells were visualised on a Nikon Eclipse 50i. Localisation analysis was performed using ImageJ (v1.50i, NIH). For each cell the outline was drawn for the cytoplasmic and nuclear regions, and for the entire cell. For each region, as well as for 5 adjacent background readings, the integrated density was measured. The localisation of the proteins is shown as the ratio between the integrated densities of the nuclear or cytoplasmic region compared to the whole-cellular ones. To assay induction of DNA damage, nuclear foci were counted from 100 transfected cells.

### Protein preparation and western blotting

To monitor association by coimmunoprecipitation, extracts were prepared by lysing cells in buffer (50mM Tris-HCl [pH7.4], 150 mM NaCl, 5% Glycerol, 1mM EDTA, 0.2% sodium deoxycholate, 1% Triton-100) containing complete protease inhibitor cocktail (Roche - catalog # 11873580001) as well as 1mM PMSF (Sigma - catalog # 93482). Immunoprecipitations were performed by incubating cleared lysates at 4°C for 4 hours using 15 μl of FLAG M2 Affinity Gel (Sigma-Aldrich - catalog #F2426) with gentle mixing. The affinity gel was then washed according to the manufacturer’s instructions. In order to minimise release of proteins bound to the gel aspecifically, immunoprecipitated proteins were eluted using 3X-FLAG peptide (Sigma - catalog # F4799) for 2 hours at 4°C according to the manufacturer’s instructions.

In order to obtain nuclear and cytoplasmic fractions, cells were incubated at 4°C for 30 minutes in hypotonic buffer A (10 mM HEPES [pH 7.6], KCl 15mM, 2mM MgCl2, 0.1mM EDTA) supplemented with protease inhibitors. After a further 5 minutes incubation on ice in presence of 1% NP40 substitute, samples centrifuged for 15’ at 1000g at 4°C. The supernatant was then cleared by centrifugation and represented the cytoplasmic fraction. After two washes with Buffer A, 1% NP40 substitute and protease inhibitors, the pellet was dissolved in Buffer B (100mM Tris HCl pH 7.4, 100mM NaCl 2mM vanadate, 2 mM EDTA, 10% Glycerol, 1% Triton X-100, 0.1% SDS) for 30 minutes on ice with mixing. The nuclei were then subjected to 8 strokes in Dounce homogenizers and cleared by centrifugation (20000g for 10 minutes at 4°C).

Protein samples were then denatured in sample buffer (0.4ml 2-Mercaptoethanol, 0.8 gr SDS, 2ml Tris HCl 1M [pH 6.8], 4ml Glycerol, 8 mg Bromophenol Blue) at 95°C for 10 minutes.

Formaldehyde crosslinking has been performed as described in Klockenbusch et al. (36). We performed crosslinking either before or after cell lysis, with overlapping results.

After SDS-PAGE, proteins were visualised by western blots using either goat anti-APOBEC1 antibody (Santa Cruz - catalog # sc-11739), rabbit anti-A1CF (Sigma - catalog # HPA037779), mouse anti-HSP90 (F-8) (Santa Cruz - catalog # sc-13119), goat anti-GAPDH V-18 (Santa Cruz - catalog # sc-20357), rabbit anti-Lamin A (H102) (Santa Cruz - catalog # sc-20680), or monoclonal anti-FLAG (Sigma - catalog # A8592). Non-conjugated primary antibodies were associated with relevant HRP-conjugated antibodies: anti-goat IgG (Santa Cruz - catalog # sc-2768), anti-rabbit IgG (Cell Signalling - catalog # 7074S), anti-Mouse IgG (Cell Signalling - catalog # 7076).

### rat APOBEC1 structure model

The rat APOBEC1 structure was modelled based on the human APOBEC1 one (pdb 6X91, (37)) using modeller 9.25 (38) using the default parameters and visualized using PyMol 2.3.4 (The PyMOL Molecular Graphics System, Schrödinger, LLC) (Supplementary File 1).

## RESULTS

### Differential inhibition of APOBEC1 activity by various catalytically inactive constructs

As titration of APOBEC1 activity using various mutants has been shown *in vitro* (30), we first aimed to test whether a dominant negative effect could be ascertained in live cells using a fluorescence-based assay previously described (33). In short, HEK293T cells are transiently transfected with APOBEC1, its RBM47 or A1CF cofactors, and the mCherry-APOB-EGFP reporter construct, which bears the editing target for APOBEC1 in frame with the chimeric coding sequence. In presence of APOBEC1-mediated RNA editing, a CAA codon in the APOB segment is edited to a Stop codon (UAA), thus leading to loss of EGFP fluorescence, which can be visualised by FACS analysis.

In order to set-up the system to titrate RNA editing using different amounts of plasmidic constructs, we first assessed the efficiency of the assay in presence of increasing amounts of competitor plasmid (Supplementary Figure 1A). Indeed, the activity of APOBEC1-mediated RNA editing decreased with the amount of transfected DNA, but the dominant negative effect of a catalytically inactive APOBEC1 (E63A) was appraisable only at lower DNA concentrations. This is probably due both to the excessive amount of DNA, which hinders the efficiency of the transfection (Supplementary Figure 1B), and to transcriptional competition among the various constructs. In order to avoid such effects, we decided to use 2 μg of cumulative APOBEC1 constructs with varying ratios of inactive/active APOBEC1 (Supplementary Figure 2). With regard to APOBEC1 cofactors RBM47 and A1CF, we tested different plasmid concentration by RNA editing assay and western blot analysis using FLAG-tagged RBM47 and A1CF. As previously noted, (39), coexpression of RBM47 gave a much stronger RNA editing signal than when A1CF was used (Supplementary Figure 3A). While it is quite possible that the specificities of the two cofactors affect the efficiency of editing, we observed that RBM47 is consistently more expressed than A1CF (Supplementary Figure 3B). In order to work with comparable amounts of protein levels, we therefore used 1 μg of plasmid for A1CF and 20ng for RBM47. Titration of APOBEC1 activity using catalytically inactive APOBEC1 (E63A) resulted in a decrease of its editing activity, an observation analogous to what described in Oka et. al (30). This effect is evident in the presence of either RBM47 (Figure 1A) or A1CF (Figure 1B). We then assessed whether two chimeras of APOBEC1 with EGFP, which we have previously shown to have different cellular localisations and activities (33), could affect the editing induced by wild-type APOBEC1. To this aim we used constructs in which catalytically inactive APOBEC1 (E63A) were fused with a non-fluorescent EGFP mutant (Y66H) (GFP-E63A and E63A-GFP in the figures) in order to avoid interference with the fluorescent reporter.

**Figure 1.**
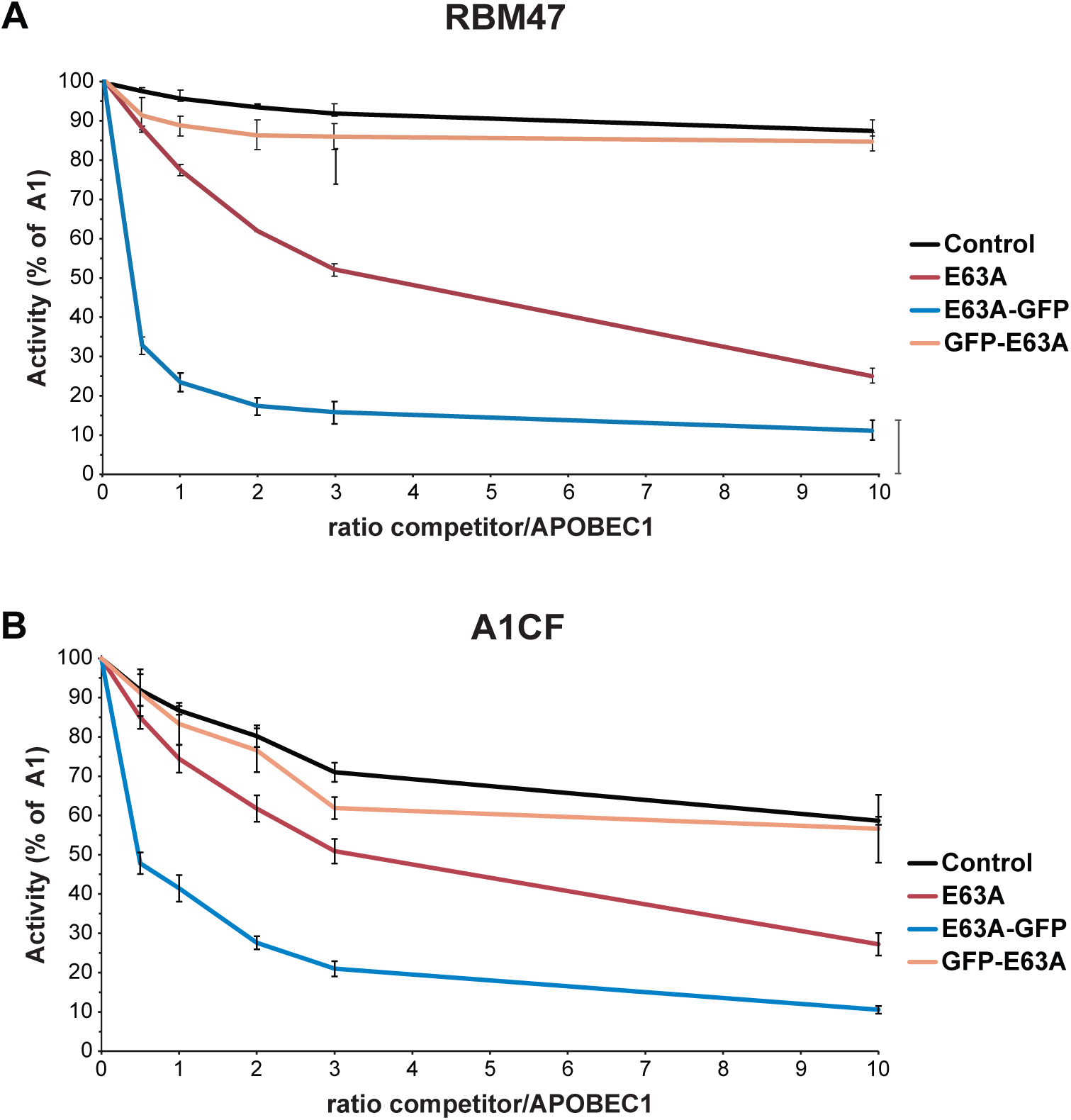
Dominant negative effects of catalytically inactive mutants on the RNA editing activity of APOBEC1. The plots represent APOBEC1-dependent RNA editing in HEK293T cells transiently cotransfected with plasmids encoding for the mCherry-APOB-EGFP reporter (1 μg) together with 2 μg of a plasmid mix expressing increasing ratios of catalytically inactive APOBEC1 mutants, either untagged or as EGFP-tagged chimeras (E63A, GFP-E63A, E63A-GFP, or a control plasmid) and wild-type APOBEC1 with 20ng of RBM47 expressing plasmid (A) or 1 μg of A1CF expressing plasmid (B). APOBEC1-dependent RNA editing of the reporter (percentage of gated cells) has been normalised to the activity of APOBEC1 in absence of competitor. The error bars represent the standard error from three experiments.

Titration with the APOBEC1^E63A^-EGFP^Y66H^ chimera (E63A-GFP) induced an even more pronounced titration of APOBEC1 activity. This can be reasoned with the localisation of E63A-GFP (33), as RNA editing takes place in the nucleus, and a nuclear mutant is probably more available to outcompete wild-type APOBEC1 in the editosome. On the other hand, inactive EGFP^Y66H^-APOBEC1^E63A^ chimera (GFP-E63A) has a freely accessible C terminus, which could support APOBEC1 dimerisation. As GFP-E63A localises mainly in the cytoplasm (33), it might interfere with APOBEC1/A1CF nuclear shuttling (28). Supposedly diminished dimerization and nuclear localisation would predict a higher inhibitory effect of GFP-E63A. Surprisingly, GFP-E63A was completely unable to titrate APOBEC1 activity. To understand these results we then analysed whether the presence of the EGFP tag alters the interplay of APOBEC1 with other components of the editosome.

### All APOBEC1 constructs interact with A1CF and RBM47

We thus assessed whether the APOBEC1-EGFP and EGFP-APOBEC1 chimeras (A1-GFP and GFP-A1) had lost the ability to interact with cofactors RBM47 or A1CF. We first used FLAG-tagged RBM47 and A1CF to immunoprecipitate the various APOBEC1 constructs from total lysates obtained from transiently cotransfected HEK293T cells (Figure 2A, B). While immunoprecipitation products were present for both APOBEC1 and the chimeras, the A1-GFP construct seemed to show the weaker interaction. In parallel, since A1CF shuttles into the nucleus in presence of APOBEC1 (28), we analysed the cellular localisation of both RBM47 and A1CF in the presence of the various APOBEC1 constructs using transiently cotransfected HEK293T cells (Figure 2C and supplementary figure 4). RBM47 is a nuclear protein by itself (25), and the presence of APOBEC1 does not induce dramatic changes in its localisation. Indeed, APOBEC1 slightly increases its nuclear accumulation, and GFP-A1 reduces it, as RBM47 interaction with the more cytoplasmic GFP-A1 probably forces some of it in the cytoplasm (Figure 2C, Supplementary figure 4). On the other hand, as expected, A1CF accumulates in the cytoplasm, and it becomes more nuclear when cells are cotransfected with APOBEC1. In a similar way, also A1-GFP increases the percentage of nuclear A1CF, albeit less than wild-type APOBEC1, in line with the weaker interaction observed in the coimmunoprecipitation experiments. In the case of GFP-A1 the accumulation of A1CF in the cytoplasm is even stronger, as expected given the cytoplasmic preference of GFP-A1 (Figure 2C. Thus, differences in the interaction of GFP-A1 with either A1CF or RBM47 cannot explain its inability to titrate APOBEC1 activity.

**Figure 2.**
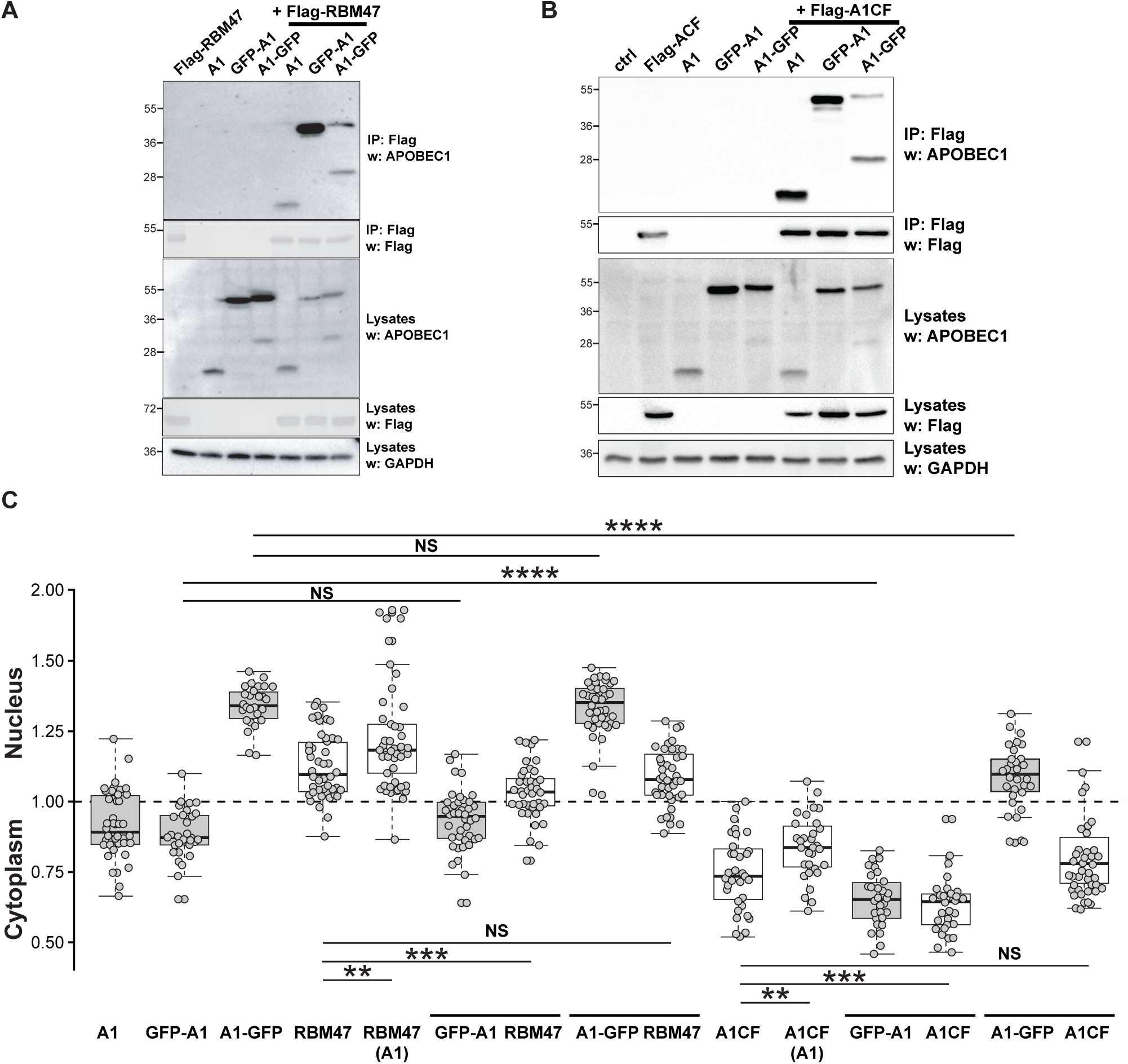
Interaction of A1CF and RBM47 cofactors with the APOBEC1 chimeras. Immunoprecipitation of FLAG-tagged RBM47 (A) or A1CF (B) brings down APOBEC1 constructs. Lysates from HEK293T cells cotransfected with the relevant FLAG-tagged cofactor together with either APOBEC1 or the GFP-tagged APOBEC1 chimeras (A1, GFP-A1, A1-GFP) were subjected to immunoprecipitation with anti-FLAG resin. Following SDS/PAGE, blots were probed with anti-APOBEC1 or anti-FLAG antibodies. Aliquots (7%) of the total-cell extracts were probed with anti-APOBEC1, anti-FLAG, or anti-GAPDH antibodies to control for expression level. The lower band (~28KDa) in the A1-GFP lane as well as the band beneath the main GFP-A1 band represent products from proteolytic cleavage of the N-terminal region of the EGFP epitope. The western blots are representative of 5 independent experiments. (C) Localisation of APOBEC1, and of RBM47 or A1CF in presence of the APOBEC1 constructs. The plot depicts the percentage of nuclear localisation of APOBEC1, RBM47 and A1CF in HEK293T cells transfected with RBM47 or A1CF alone or in combination with APOBEC1 constructs (A1, GFP-A1, A1-GFP). Thirty cells per sample were analysed, median, 25^th^/75^th^ percentiles and 25^th^/75^th^±1.5IQR are indicated in the boxplot. The levels of statistical significance (one-way ANOVA coupled with Tukey’s test) are indicated: *, P<0.02; **, P<0.01; ***, P<0.001; ****, P<0.0001). Representative images of HEK293T cells transfected with either RBM47 or A1CF alone or in combination with the APOBEC1 constructs (A1, GFP-A1, A1-GFP) are shown in Supplementary figure 4.

### APOBEC1-EGFP does not dimerise

Considering the correlation between APOBEC1 editing ability and its dimerisation, even though its meaning is controversial (30-32), we wondered whether interaction of the APOBEC1 chimeras with APOBEC1 could explain the results of the titration experiments. We tested the ability of APOBEC1 to form heterodimers with the chimeras in transiently transfected HEK293T cells: coimmunoprecipitation of either wild-type APOBEC1, GFP-A1, or A1-GFP using FLAG-tagged APOBEC1 reveals that the only construct with a very limited ability to form heterodimers is A1-GFP (Figure 3A).

**Figure 3.**
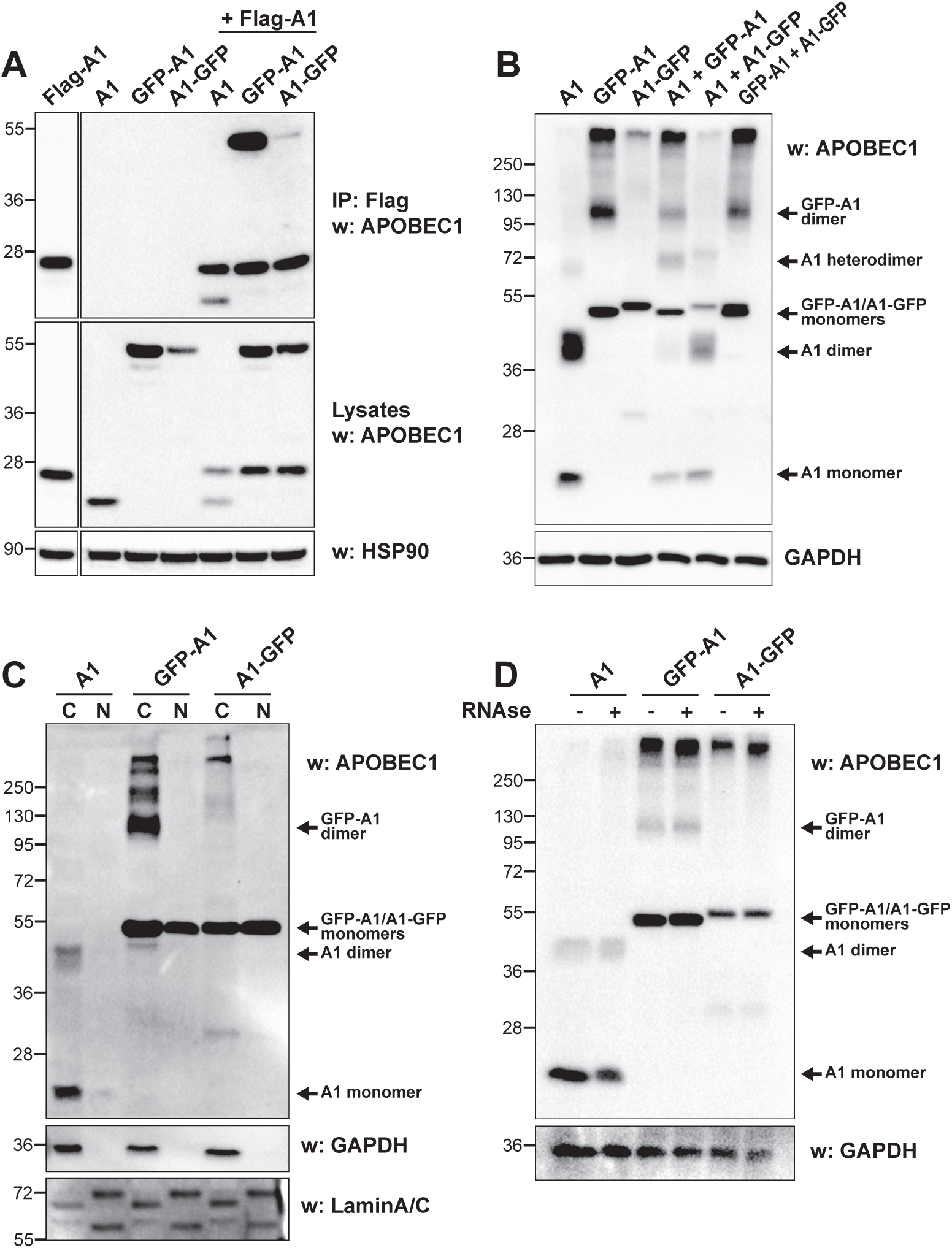
The APOBEC1-EGFP chimera displays reduced dimerisation. (A) Immunoprecipitation of FLAG-tagged APOBEC1 shows a decreased dimerisation of the APOBEC1-EGFP chimera. Lysates from HEK293T cells cotransfected with FLAG-tagged APOBEC1 together with either APOBEC1 or the EGFP-tagged APOBEC1 chimeras (A1, GFP-A1, A1-GFP) were subjected to immunoprecipitation with anti-FLAG resin. Following SDS/PAGE, western blots were probed with anti-APOBEC1 antibodies. Aliquots (7%) of the total-cell extracts were probed with anti-APOBEC1 or anti-HSP90 antibodies to control for expression level. (B) Crosslinking of the APOBEC1 constructs in whole cell lysates, (C) in cytoplasmic/nuclear fractions, or (D) after treatment with RNAse A. HEK293T cells were transiently cotransfected as indicated and were treated with 2% formaldehyde (10’ at room temperature). The crosslinking was blocked with 1.25M glycine and cells were lysed. Following SDS/PAGE, western blots were probed with anti-APOBEC1 antibodies. All western blots are representative of at least three independent experiments.

We next confirmed the inability of A1-GFP to dimerise by formaldehyde cross-linking in transiently transfected HEK293T cells. After treatment with formaldehyde, cells were lysed, and western blotting was performed to visualise APOBEC1 (Figure 3B). A lower band for the wild-type APOBEC1 at the expected molecular weight for monomeric APOBEC1 (27kDa) was associated to a stronger one, presumably the dimeric form (expected molecular weight 54kDa, but crosslinks alter the apparent molecular weight). Other weaker bands were present as well (overexposed image in Supplementary Figure 4A), possibly representing other aggregation forms. An analogous pattern was apparent for the GFP-A1 chimera (56kDa for the monomer; 112kDa for the dimeric form), suggesting that both APOBEC1 and the GFP-A1 chimera are able to dimerise. On the other hand, only one band corresponding to the monomer was present for the A1-GFP chimera. Even though a much weaker band of high molecular weight was present (overexposed image in Supplementary Figure 5A), these results corroborate the inability of the A1-GFP chimera to dimerise. Coexpression of APOBEC1 together with the chimeras leads to the formation of bands likely representing heterodimers (83 kDa, Figure 3B). Yet, the ratio between the homodimeric APOBEC1 band and the heterodimeric one suggests that the interaction between APOBEC1 and A1-GFP is much weaker than the one between APOBEC1 and GFP-A1. Coexpression of both chimeras together result in a pattern identical to that of GFP-A1 alone.

Crosslinking experiments after nuclear/cytoplasmic fractionation (Figure 3C; Supplementary Figure 5B) reveal that the dimerisation bands of GFP-A1 disappear completely in the nuclear fraction, thus leaving only the monomeric form in the nucleus. While this differential localisation of the GFP-A1 dimeric form could be due to its size being large enough to prevent its nuclear import (albeit a decreased nuclear localisation of the APOBEC1 dimer might be present as well; Supplementary Figure 5B), this reinforces the notion that dimerisation of APOBEC1 is not needed for its activity. On the other hand, treatment with RNAse does not affect dimerisation (Figure 3D), suggesting that, as in the case of human APOBEC1 (37), the dimerisation depends on protein-protein interaction. In all samples where the chimeras were expressed a very high molecular weight band was present as well: considering its absence in the sample where APOBEC1 alone is expressed, this band might be due to the presence of EGFP-induced aggregation in the cytoplasm.

### Dimerisation of APOBEC1 is dispensable for RNA editing

Considering that the A1-GFP was the only construct that, while maintaining its ability to trigger RNA editing, had a different behaviour with regard to interacting partners -in particular its inability to dimerise-we wondered how its activity on the RNA editing target would change in presence of the inactive APOBEC1 constructs. We thus titrated the activity of an APOBEC1-EGFP^Y66H^ (A1-GFP) construct using the catalytically inactive ones. As expected, the E63A-GFP chimera is able to titrate A1-GFP activity (Figure 4) in a similar manner to what observed with the titration of wild-type APOBEC1 using either the inactive E63A mutant or the E63A-GFP chimera (Figure 1). Similarly, the GFP-E63A mutant is unable to outcompete the activity of A1-GFP. What is drastically different is the RNA editing activity in presence of the E63A mutant, which is now unable to titrate the activity of A1-GFP. On one hand, the lack of competition by E63A on the dimerisation-unable A1-GFP proves that dimerisation of APOBEC1 is not necessary for its RNA editing. On the other one, especially in presence of the A1CF cofactor, the activity observed is even higher than that observed in presence of the negative control plasmid. This might suggest that some other factor affects the activity of APOBEC1 in the nucleus.

**Figure 4.**
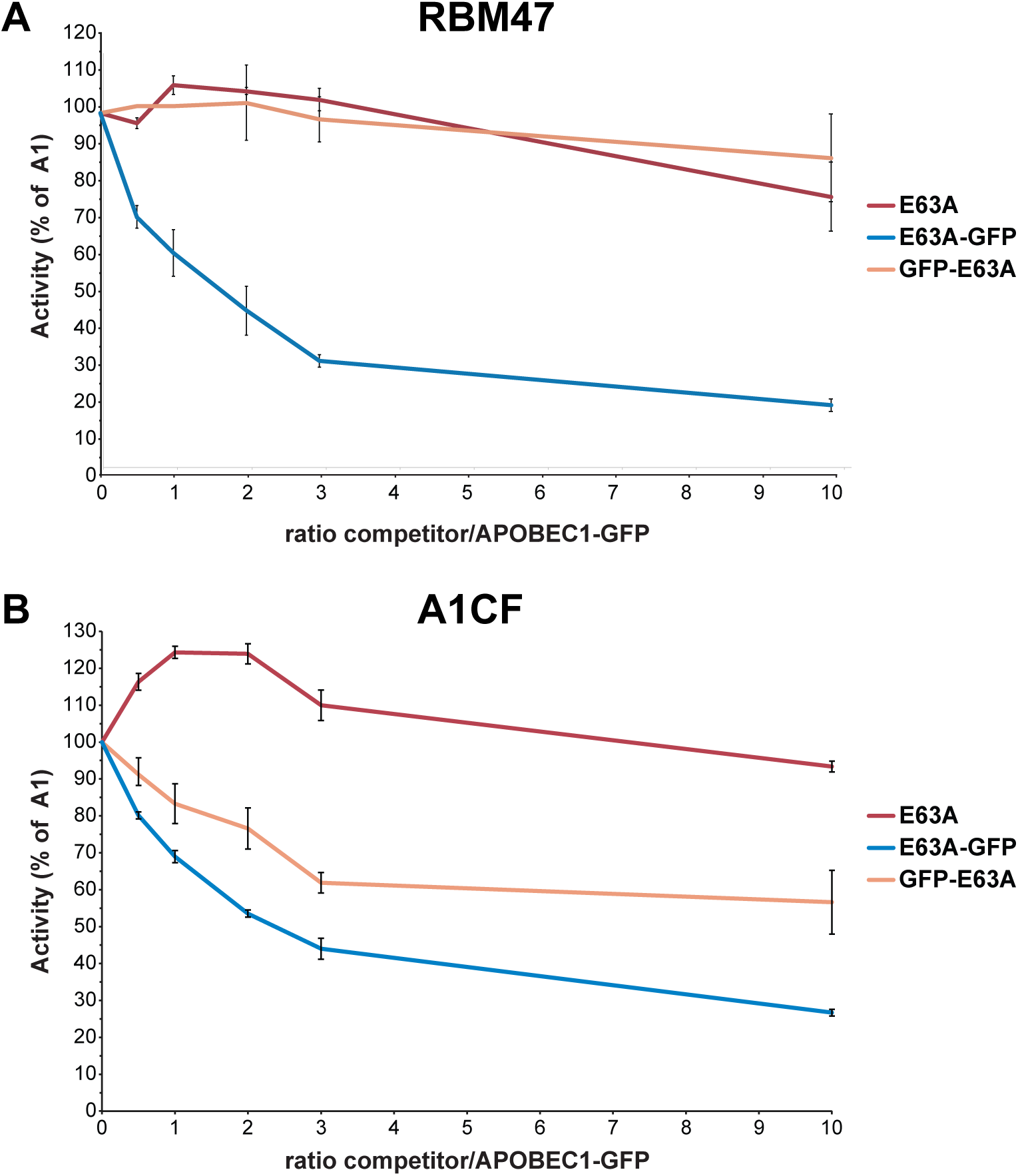
Titration of RNA editing activity mediated by the monomeric EGFP-tagged APOBEC1. The plots represent APOBEC1-dependent RNA editing in HEK293T cells transiently cotransfected with plasmids encoding for the mCherry-APOB-EGFP reporter (1 μg) together with 2 μg of a plasmid mix expressing increasing ratios of catalytically inactive APOBEC1 constructs (E63A, GFP-E63A, E63A-GFP) and the APOBEC1-EGFP chimera with 20ng of RBM47 expressing plasmid (A) or 1 μg of A1CF expressing plasmid (B). APOBEC1-dependent RNA editing of the reporter (percentage of gated cells) has been normalised to the activity of APOBEC1-EGFP in absence of competitor. The dotted lines report the RNA editing activity shown in Figure 1. The error bars represent the standard deviation from three experiments. The experiments shown in Figure 1 and Figure 4 were performed concurrently.

A recently described structure of human APOBEC1 allowed the identification of a single point mutant (L173Q) unable to form dimers *in vitro* (37). Based on that work, we tested the effects of a L173Q mutation on rat APOBEC1. Intriguingly, the L173Q mutant is still able to form dimers in cells (Supplementary Figure 6). We therefore modelled the structure of rat APOBEC1 based on that of human APOBEC1 and identified position G227 as a potential site at the interface of the dimer (Figure 5). While the G227A mutant was still able to form homodimers, as evidenced by formaldehyde crosslinking and co-immunoprecipitation (Figure 6A and B), a double mutant L173A/G227A was not. Yet, the L173A/G227A mutant retains the ability to interact with wild-type APOBEC1 (Figure 6A and B). As these mutants maintain the ability to interact with both RBM47 and A1CF (Figure 6C), we tested the activity of the L173A/G227A mutant in the cellular assay, where its activity surpasses that of both wild-type APOBEC1 and A1-GFP chimera (Figure 7A and B).

**Figure 5.**
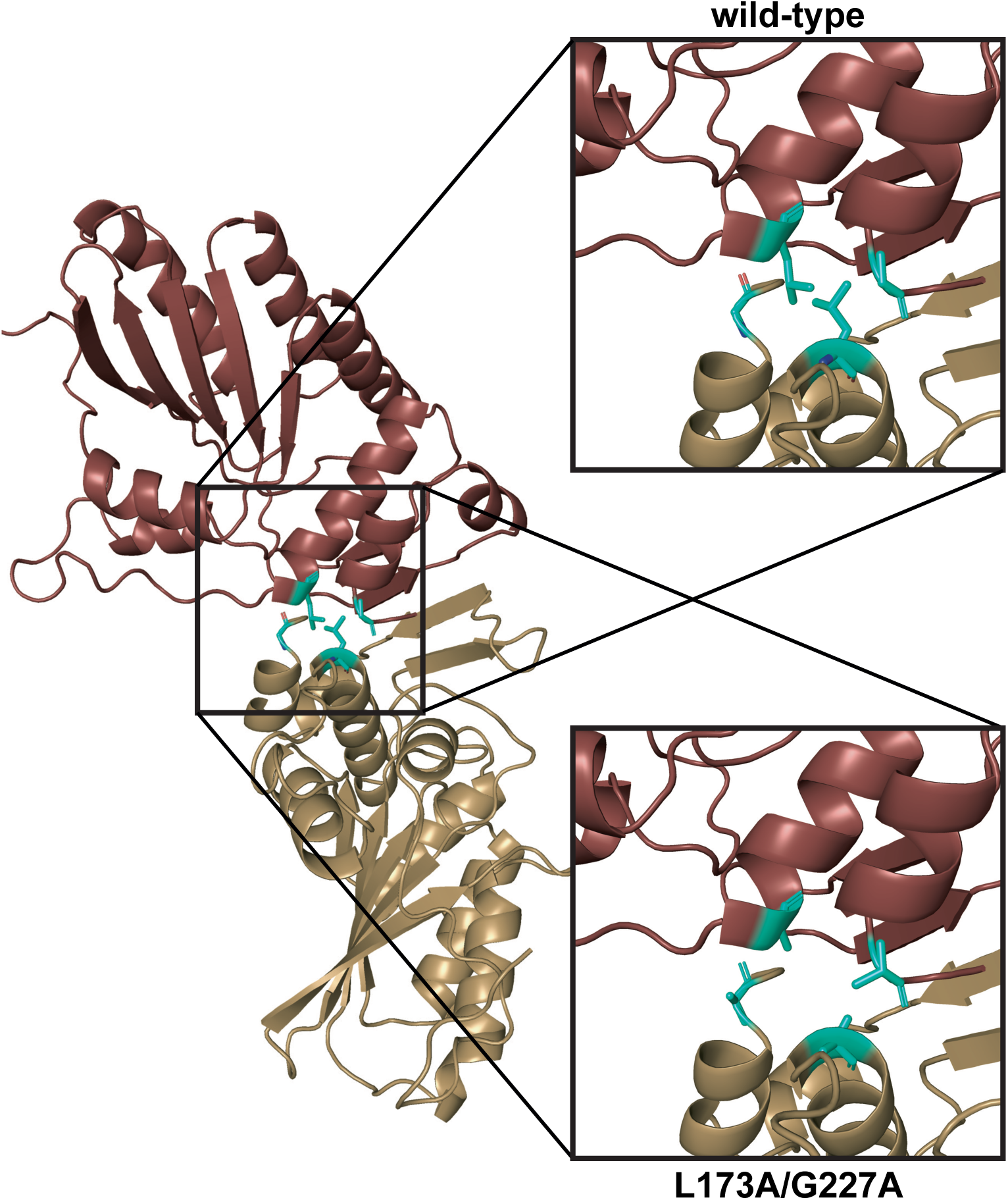
Ribbon view of the rat APOBEC1 dimer modelled on the human APOBEC1 dimer structure (37). The 173 and 227 residues are highlighted in green. The upper panel depicts the wildtype residues, which are conserved between human and rat sequence. The lower panel depicts the L173A and the G227A mutations.

**Figure 6.**
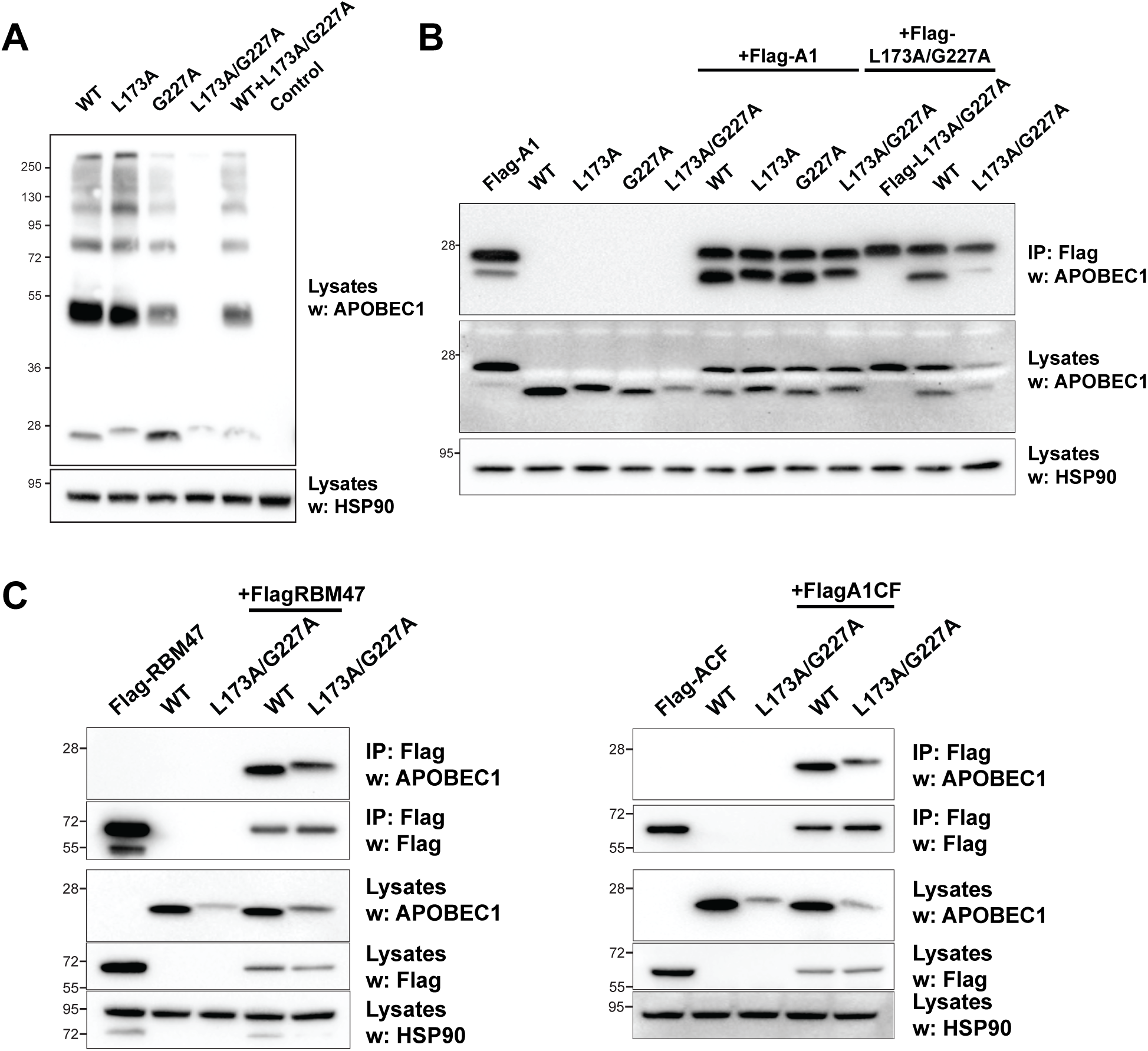
The APOBEC1^L173A/G227A^ mutant displays reduced self-dimerisation but retains the interaction with its cofactors. (A) Crosslinking of the APOBEC1 constructs in whole cell lysates. HEK293T cells were transiently cotransfected as indicated and were treated with 2% formaldehyde (10’ at room temperature). The crosslinking was blocked with 1.25M glycine and cells were then lysed. Following SDS/PAGE, western blots were probed with anti-APOBEC1 antibodies. (B) Coimmunoprecipitation of the APOBEC1 mutants using FLAG-tagged APOBEC1, wild-type (Flag-A1) or the APOBEC1^L173A/G227A^ mutant (Flag-L173A/G227A). Lysates from HEK293T cells that had been cotransfected with FLAG-tagged APOBEC1 constructs together with either APOBEC1 or APOBEC1 mutants (L173A, G227A or L173A/G227A) were subjected to immunoprecipitation using anti-FLAG resin. Following SDS/PAGE, western blots were probed with anti-APOBEC1 antibodies. Aliquots (7%) of the total-cell extracts were probed with anti-APOBEC1, anti-FLAG, or anti-HSP90 antibodies to control for expression level. C) Interaction of A1CF and RBM47 cofactors with the APOBEC1 mutants. Immunoprecipitation of FLAG-tagged RBM47 (left) or A1CF (right) brings down the APOBEC1 mutants. Lysates from HEK293T cells cotransfected with the relevant FLAG-tagged cofactor together with either APOBEC1 or the APOBEC1 mutants (L173A, G227A or L173A/G227A) were subjected to immunoprecipitation with anti-FLAG resin. All western blots are representative of at least three independent experiments.

**Figure 7.**
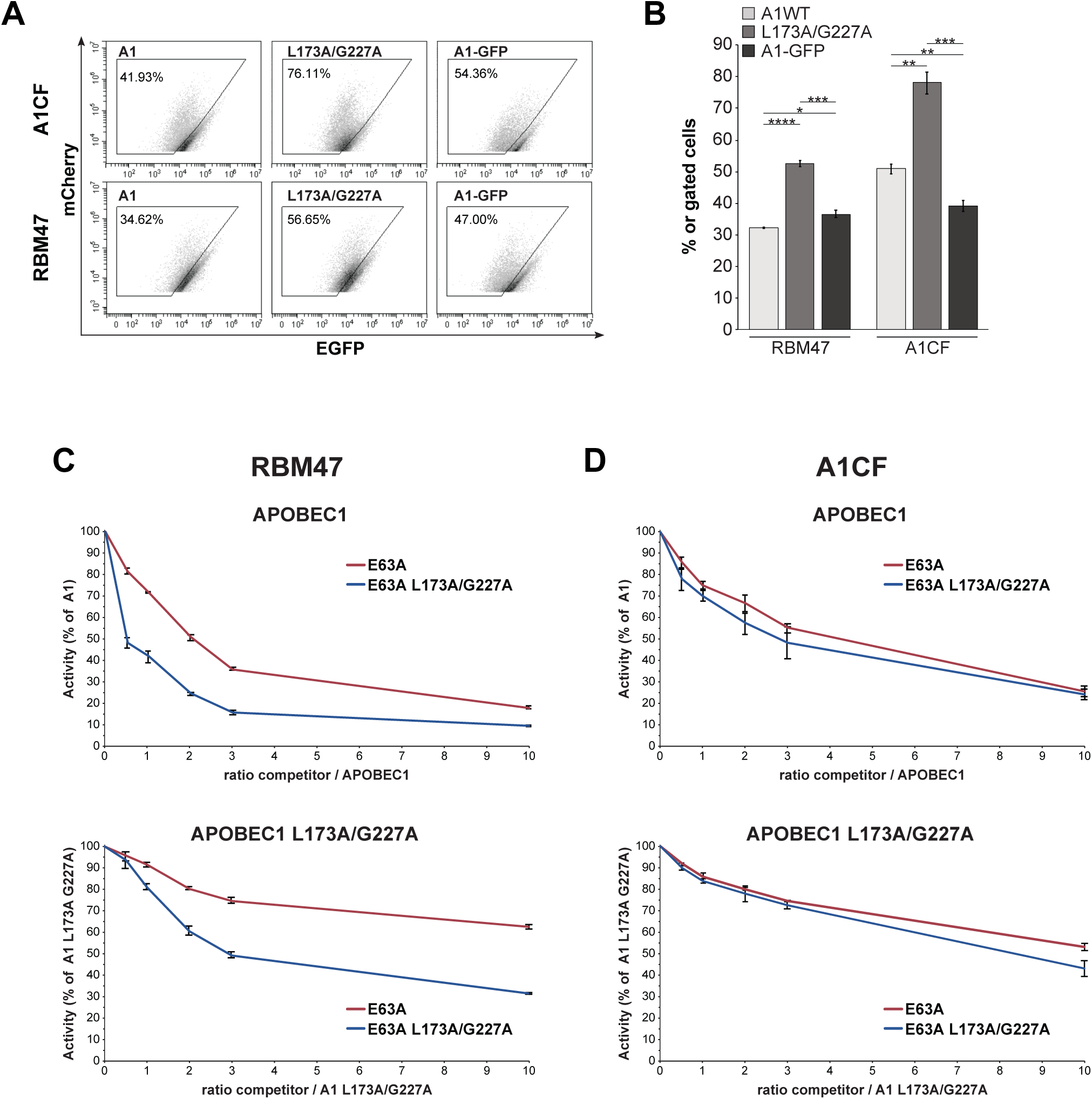
RNA editing of the APOBEC1^L173A/G227A^ mutant. (A) Representative FACS analysis and (B) bar graph showing APOBEC1-dependent RNA editing in HEK293T cells transiently cotransfected with plasmids encoding for the mCherry-APOB-EGFP reporter (1 μg), the A1CF (1 μg) or RBM47 (20ng) cofactors, and either wild type APOBEC1 (A1), the mutant (L173A/G227A) or the chimeric APOBEC1 (A1-GFP). The error bars represent the standard deviation from three experiments. The levels of statistical significance (one-way ANOVA coupled with Tukey’s test) are indicated: *, P<0.02; **, P<0.01; ***, P<0.001; ****, P<0.0001). Titration of APOBEC1-dependent RNA editing in presence of A1CF (C) or RBM47 (D). Cotransfections were carried out as described above but, in lieu of APOBEC1, a plasmid mix (2 μg) with increasing ratios of catalytically inactive APOBEC1 constructs (E63A or E63A/L173A/G227A) and either APOBEC1 or APOBEC1^L173A/G227A^ mutant was used APOBEC1-dependent RNA editing of the reporter (percentage of gated cells) has been normalised to the activity of APOBEC1 or APOBEC1^L173A/G227A^ in absence of competitor.

We then performed experiments to titrate the activity of either APOBEC1 or APOBEC1^L173A/G227A^ with catalytically inactive mutants (E63A; E63A/L173A/G227A). Indeed, when RBM47 was used as cofactor (Figure 7C), the constructs bearing L173A/G227A mutations behaved analogously to the APOBEC1-EGFP chimeras (E63A-GFP and A1-GFP, Figures 1 and 4). When A1CF was used, the differences were substantially lower (Figure 7D). At this time, we cannot explain the difference in titration in presence of A1CF and RBM47. It might be due to intrinsic differences between the two cofactors (e.g., their localisation). Nonetheless, the increased activity of the L173A/G227A mutant and its increased titration of APOBEC1 suggest that dimerisation of APOBEC1 might have a regulatory function on RNA editing.

### APOBEC1 dimerisation limits its degradation

The lower expression levels of both A1-GFP and L173A/G227A compared to the other constructs (lysates in Figure 3A and Figure 6B) could be an indication of the instability of monomeric APOBEC1 in the cell. Indeed, treatment of transiently transfected cells with the protease inhibitor MG132 induced an increase of both A1-GFP and L173A/G227A (Figure 8A and B), suggesting that monomeric APOBEC1 is actively degraded through the proteasome.

**Figure 8.**
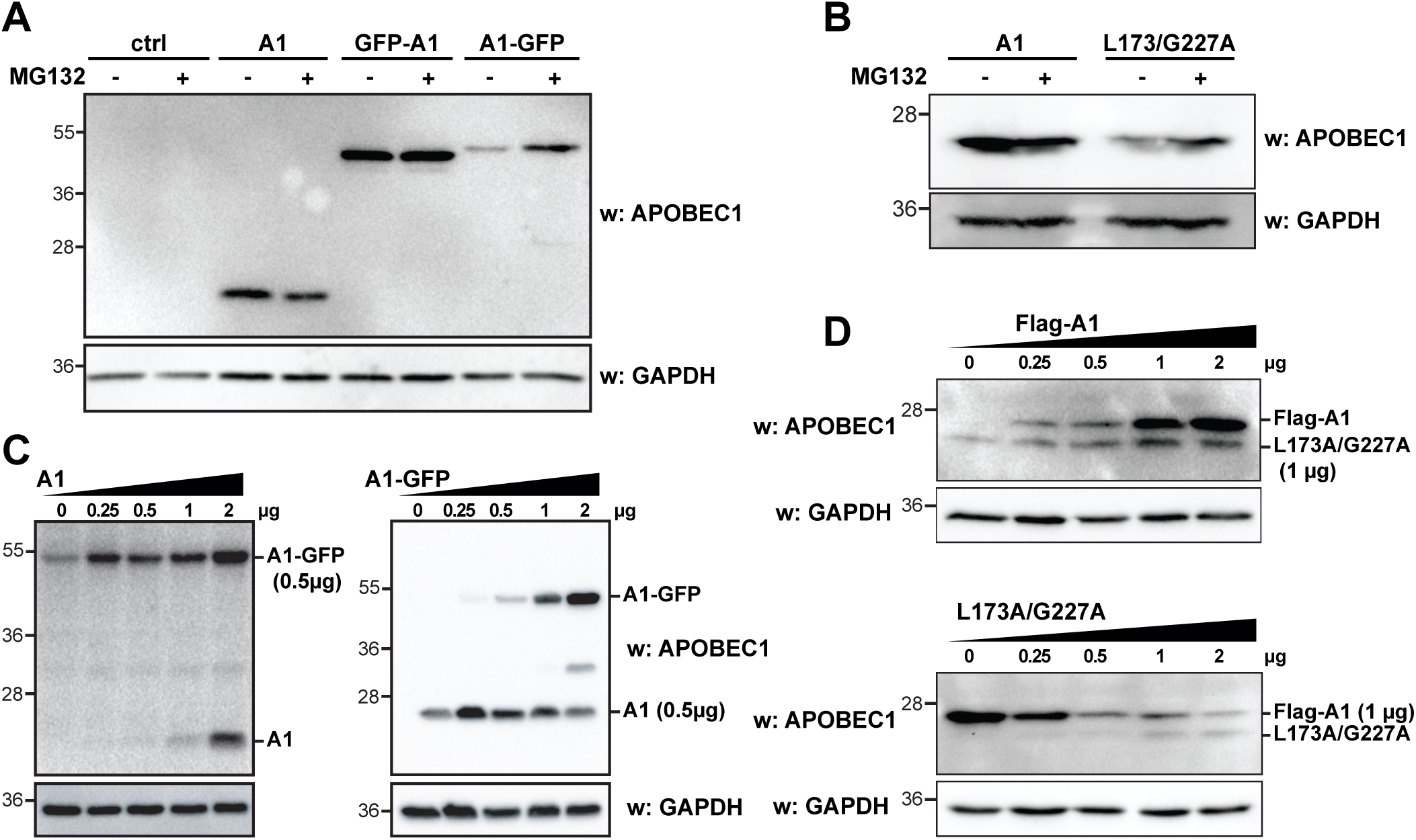
Dimerisation and degradation of APOBEC1. Representative western blots showing the expression levels of APOBEC1, the EGFP-tagged chimeras (A), or the APOBEC1^L173A/G227A^ mutant (B) after treatment with proteasome inhibitor MG132. Representative western blots showing the expression levels of fixed amounts (0.5 μg) of either APOBEC1-EGFP or APOBEC1 (C) and FLAG-tagged APOBEC1 or APOBEC1^L173A/G227A^ (D) in presence of increasing amounts of the other one as indicated. HEK293T cells were transiently co-transfected with the indicated plasmids and lysed after 36 hours. All western blots are representative of at least three independent experiments. GAPDH has been used to control for expression level.

Taking together the increased activity of the monomeric A1-GFP in presence of increasing amounts of dimeric E63A (Figure 4) with the increased expression of both A1-GFP and L173A/G227A in presence of A1 (lysates in Figure 3A and Figure 6B), we hypothesized that dimerisation might protect APOBEC1 from degradation. In order to test this, we cotransfected fixed amounts of A1-GFP or L173A/G227A with increasing amounts of APOBEC1 or Flag-tagged APOBEC1, respectively (in order to allow discrimination by western blot analysis; Figure 8C and D). In line with our hypothesis, the levels of monomeric APOBEC1 forms (A1-GFP or L173A/G227A) increased when increasing amounts of dimeric APOBEC1 were used (A1 or Flag-A1). Conversely, either no increase or decrease of APOBEC1 levels were perceptible when APOBEC1 constructs that could form dimers were cotransfected with increasing amounts of A1-GFP or L173A/G227A, respectively. This suggests that the increased activity we observed with the titration of A1-GFP and L173A/G227A might be the result of their increased availability in conditions in which a heterodimer could form.

### APOBEC1 dimerisation titrates its DNA targeting ability

Since APOBEC1 can act on DNA (5, 6, 37, 40), we then tested whether dimerisation could affect the ability of APOBEC1 to target DNA. We assessed the induction of DNA damage by γ-H2AX foci formation. Despite reduced expression of the L173A/G227A mutant (Figure 6), its presence in transiently transfected HEK293T cells leads to increased DNA damage compared to wild-type APOBEC1 with regard both to the number of DNA repair foci per nucleus and to the percentage of cells bearing DNA damage (Figure 9A, B). We then tested the effects of a mix of active/inactive APOBEC1 constructs to test whether the inactive mutants could protect the active forms from degradation and increase their mutagenicity. As expected, the presence of catalytically inactive APOBEC1 (E63A) led to an increase of the γ-H2AX (+) population. When inactive E63A/L173A/G227A mutant was cotransfected with the monomeric L173A/G227A the mutagenic activity increased even more (Figure 9C and supplementary figure 7).

**Figure 9.**
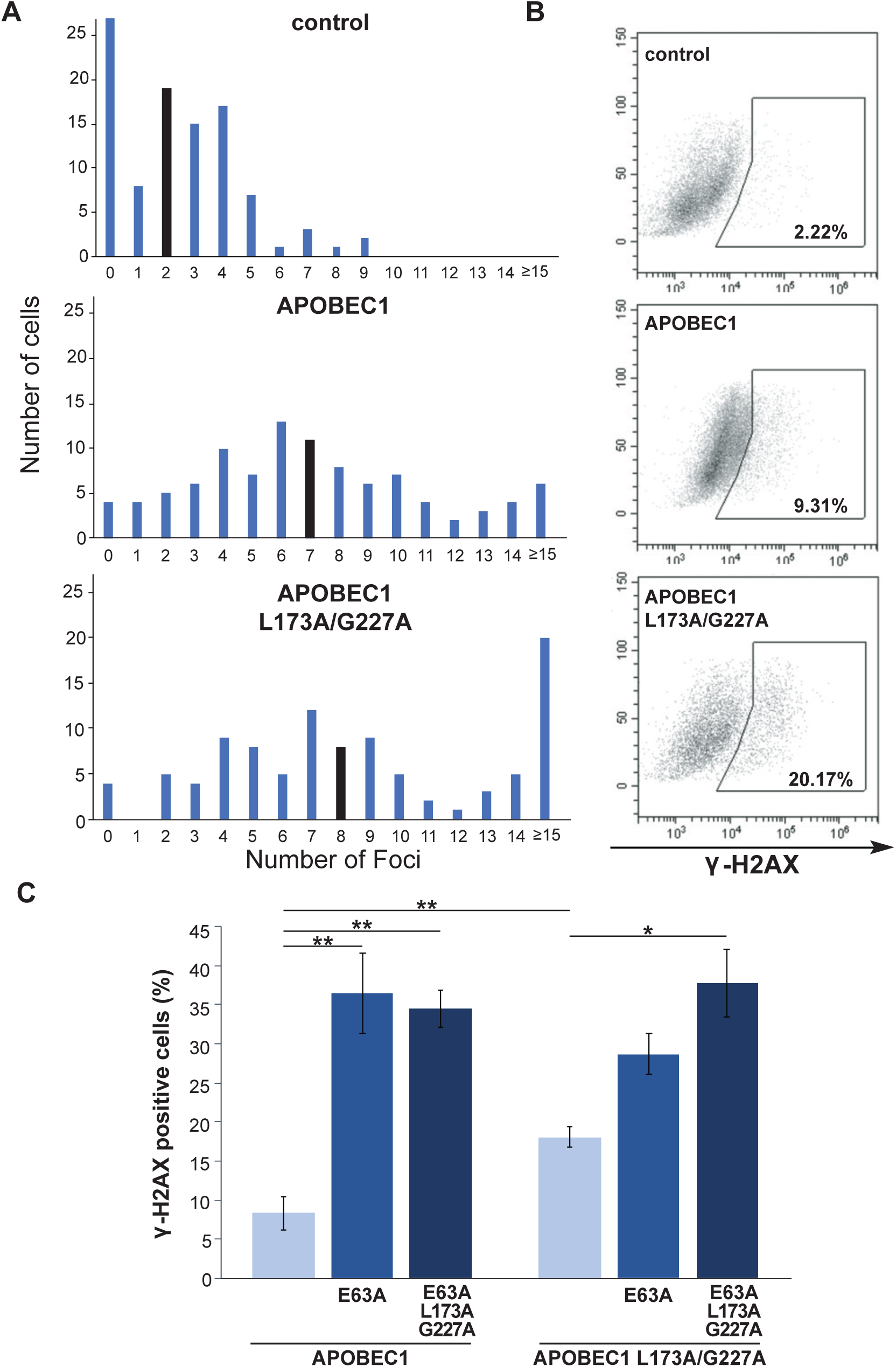
APOBEC1 induction of DNA damage. (A) Bar plot showing the number of DNA damage foci induced after transient transfection with a control plasmid, wild-type APOBEC1 or the APOBEC1^L173A/G227A^ mutant. 48 hours after transfection, cells were fixed and immunofluorescence analysis using anti-γ-H2AX antibody was performed. 100 cells per sample were analysed. The median number of foci per nucleus is indicated as a black bar. (B) Representative FACS profiles of transfected cells after intracellular staining with anti-γ-H2AX antibody. (C) Induction of DNA damage in HEK293T cells transfected with either APOBEC1 (A1) or APOBEC1^L173A/G227A^ (L173A/G227A) and an excess of catalytically inactive APOBEC1 (E63A or E63/L173A/G227A). Cells were transfected with a plasmid mix containing catalytically active and inactive APOBEC1 constructs in a 1:3 ratio (2 μg). The levels of statistical significance (one-way ANOVA coupled with Tukey’s test) are indicated: *, P<0.02; **, P<0.01; ***, P<0.001; ****, P<0.0001). In all experiments shown a cotransfected EGFP expressing plasmid (100ng) was used to identify transfected cells. Representative FACS profiles are shown in Supplementary Figure 7.

## DISCUSSION

Dissection of the mode of action of the AID/APOBECs has always been a central research theme, as their ability to target DNA and RNA can have profound repercussions on the stability of both genetic and transcribed information in the cell.

It is important to discriminate two steps in the mechanism of action of the AID/APOBECs: the targeting of the AID/APOBEC to the nucleic acid in a specific cellular compartment, and its ability to act on the target itself. A correlation between dimer formation and association with the DNA substrate has been observed through several approaches, sometimes with conflicting interpretations (40-52). Structural analysis suggests that dimerisation of individual APOBEC proteins might be either an intrinsic property of the AID/APOBECs (37, 47, 53-55) or mediated by nucleic acids (56-59). Yet monomeric APOBECs can target ssDNA and are active in cells and *in vitro* (43, 45, 55, 60-63). While most of these studies address the importance of dimer formation towards viral or mobile element restriction, for which binding to the RNA is relevant for the packaging into the virion, only little is known about targeting specific RNA transcripts: it has been hypothesised that APOBEC1 acts as a dimer within the editing complex (31), yet the use of APOBEC1 mutants lacking the ability to dimerise led to contradictory results (30, 32).

In this context, our results allow for the first time the definition of a model where dimerisation and degradation are strictly connected in regulating APOBEC1 activity (Figure 10). Our data indicate that APOBEC1 ability to dimerise is dispensable for its RNA editing activity and its monomeric forms are more active. Moreover, the increased levels of the monomeric constructs (A1-GFP and L173A/G227A) either after treatment with the MG-132 proteasomal inhibitor (Figure 8A, B) or through coexpression of APOBEC1 (Figure 8C, D) suggest the existence of a correlation between dimerisation of APOBEC1 and its degradation. Thus, increased activity can be obtained when a monomeric APOBEC1 is used, but at the price of increased degradation (Figure 10, middle). On the other hand, imbalances in dimer formation can lead to heightened activity with reduced degradation (Figure 10, bottom).

**Figure 10.**
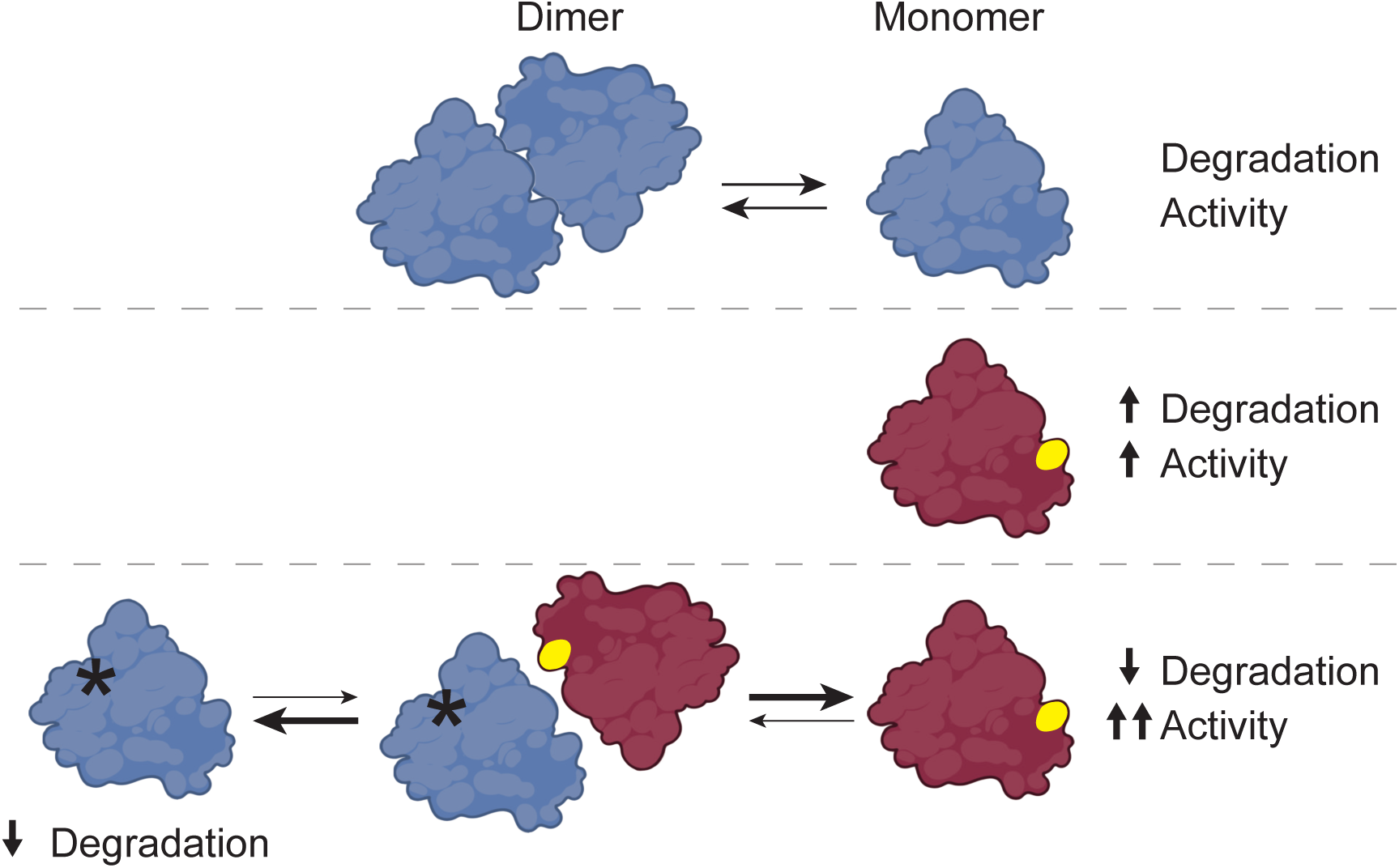
Schematic model of the effects of dimerization capability on the editing activity and the degradation of APOBEC1. The dimerization-capable wild-type APOBEC1 is shown in blue, the monomeric APOBEC1 L173A G227A mutant in red. The catalytic inactive APOBEC1 E63A is labelled with a star.

With regard to the interplay of APOBEC1 with its cofactors, we observe a weaker interaction of the APOBEC1-EGFP chimera with both RBM47 and A1CF. Analogously, the APOBEC1-EGFP construct induces only a moderate shift in A1CF localisation compared to APOBEC1, while retaining its activity on the APOB target. This suggests that even a fleeting interaction with A1CF is sufficient to allow significant editing of the transcript by APOBEC1. As it has been observed that a weaker interaction between APOBEC1 and A1CF can modulate hyperediting of the APOB transcript (64), it will be interesting to assess in a cellular model less reliant on overexpression whether APOBEC1 dimerisation affects hyperediting as well.

Similar to the other AID/APOBECs, APOBEC1 is also able to act on DNA (5, 65–70), and its mutagenic effect has been linked to cancer (5, 6). Even with regard to APOBEC1 mutagenic activity, its dimerisation dynamics are relevant, as we observe increased induction of DNA damage when monomeric APOBEC1 is expressed (Figure 9A and B). In line with our model (Figure 10), an excess of inactive APOBEC1, whose monomeric component outcompetes active APOBEC1 for degradation, leads to increased activity on DNA (Figure 9C). Even when the monomeric APOBEC1 (L173A/G227A) is considered, its maximum activity is achieved only when an excess of its corresponding inactive form is present. It becomes evident the similarity with Activation Induced Deaminase (AID), the family member that triggers all antigen-driven antibody diversification processes, which is predominantly degraded in the nucleus through the proteasome (71). Since the monomeric and dimeric forms of AID interact with different molecules (72), it could be possible that APOBEC1 monomers are preferentially degraded through interaction with another cellular factor, be it nucleic acids or proteins.

Whatever the mechanisms, the functional meaning of APOBEC1 dimerisation seems to lie in an effort to reduce its genotoxicity. It will be interesting to see whether dimerisation and degradation represent regulatory layers also for the APOBEC3s, in particular for those linked to the onset of the APOBEC mutational signature in the cancer genomes. As we used mutants to investigate dimerisation, we limited our focus to the determinants on APOBEC1. It is evident that a cellular machinery is in place to regulate APOBEC activity, and -in absence of alterations in the APOBEC proteins-it is possible that their oncogenic potential might arise from a failure in their regulation.

## DATA AVAILABILITY

Plasmids are available through the Authors or through Addgene (https://www.addgene.org): pEGFP ratAPOBEC1-EGFP, #112857; pEGFP ratA1, #112858; pEGFP mCherry-apoB-EGFP, #112859; pEGFP human A1CF, #112860; pEGFP ratA1 L173A/G227A #167169. Flow cytometry data is available on the FlowRepository (ID: FR-FCM_Z3H3).

## SUPPLEMENTARY DATA

Supplementary Data are available at NAR online.

## SUPPLEMENTARY MATERIAL

**Supplementary Figure 1.** Linearity of APOBEC1 activity with increasing amount of plasmid DNA RNA editing of rat APOBEC1 in HEK293T cells transiently cotransfected with plasmids encoding for the mCherry-APOB-EGFP reporter (1 μg), APOBEC1 (0.5 μg), A1CF (1 μg), and increasing amounts of either a control plasmid (ctrl) or a catalytically inactive APOBEC1 mutant (E63A). (A) the panel shows the APOBEC1-dependent RNA editing of the reporter (percentage of gated cells) normalised to the activity of APOBEC1 in absence of competitor DNA. The ratio of the competitor DNA with APOBEC1 is shown in red, and the amount of total DNA used in the transfection is shown in parentheses. (B) The lower panel shows the transfection efficiency (mCherry(+) cells) for each point. The error bars represent the standard deviation from three experiments.

**Supplementary Figure 2.** Coexpression of different ratios of APOBEC1 constructs. Representative western blots showing the expression levels of various combinations of APOBEC1 constructs in transiently transfected HEK293T cells. 2 μg of total plasmid DNA were transfected in cells using the ratios indicated on the blots. There is not representative blot for GFP-A1/A1-GFP cotransfections, as the identical molecular weight does not allow their discrimination.

**Supplementary Figure 3.** (A) RNA Editing Activity of APOBEC1 in presence of either RBM47 or A1CF and RBM47 cofactors. Increasing amounts (20 ng - 1 μg) of A1CF (blue) and RBM47 (red) were cotransfected in HEK293T cells together with 1 μg of editing reporter plasmid and 1 μg of APOBEC1. (B) Comparison of the expression levels of the cofactors. Increasing amounts (20 ng - 1 μg) of FLAG-tagged A1CF and RBM47 were cotransfected in HEK293T cells and their expression was assayed by western blot using anti-FLAG antibody.

**Supplementary Figure 4.** Localisation of APOBEC1 (A), and of RBM47 (B) or A1CF (C) in presence of the APOBEC1 constructs. Representative images of HEK293T cells transfected with either RBM47 or A1CF alone or in combination with the APOBEC1 constructs (A1, GFP-A1, A1-GFP). An empty plasmid was used as a negative control. Localisation of the various proteins was assessed by GFP fluorescence for the GFP-tagged APOBEC1 chimeras, and by anti-FLAG or anti-A1CF antibodies for visualization of APOBEC1, RBM47 or A1CF, respectively.

**Supplementary Figure 5.** Different exposures of the crosslinking experiments shown in figure 3B (A) and C (B).

**Supplementary Figure 6.** Crosslinking of APOBEC1 L173Q in whole cell lysates. HEK293T cells were transiently cotransfected as indicated and were treated with 2% formaldehyde (10’ at room temperature). The crosslinking was blocked with 1.25M glycine and cells were then lysed. Following SDS/PAGE, western blots were probed with anti-APOBEC1 antibodies.

**Supplementary Figure 7.** Representative FACS profiles of transfected cells after intracellular staining with anti-γ-H2AX antibody. Induction of DNA damage in HEK293T cells transfected with either APOBEC1 (A1) or APOBEC1^L173A/G227A^ (L173A/G227A) and an excess of catalytically inactive APOBEC1 (E63A or E63/L173A/G227A). Cells were transfected with a plasmid mix containing catalytically active and inactive APOBEC1 constructs in a 1:3 ratio (2 μg). In all experiments shown a cotransfected EGFP expressing plasmid (100ng) was used to identify transfected cells.

**Supplementary File 1.** Pymol workspace file with the structure of the modelled rat APOBEC1.

## REFERENCES

1. Salter, J.D., Bennett, R.P. and Smith, H.C. (2016) The APOBEC Protein Family: United by Structure, Divergent in Function. Trends Biochem Sci 41, 578–594.

2. Swanton, C., McGranahan, N., Starrett, G.J. and Harris, R.S. (2015) APOBEC Enzymes: Mutagenic Fuel for Cancer Evolution and Heterogeneity. Cancer Discov 5, 704–712.

3. Yamanaka, S., Balestra, M.E., Ferrell, L.D., Fan, J., Arnold, K.S., Taylor, S., Taylor, J.M. and Innerarity, T.L. (1995) Apolipoprotein B mRNA-editing protein induces hepatocellular carcinoma and dysplasia in transgenic animals. Proc Natl Acad Sci U S A 92, 8483–8487.

4. Blanc, V., Henderson, J.O., Newberry, R.D., Xie, Y., Cho, S.-J., Newberry, E.P., Kennedy, S., Rubin, D.C., Wang, H.L., Luo, J. and Davidson, N.O. (2007) Deletion of the AU-rich RNA binding protein Apobec-1 reduces intestinal tumor burden in Apc(min) mice. Cancer Res 67, 8565–8573.

5. Harris, R.S., Petersen-Mahrt, S.K. and Neuberger, M.S. (2002) RNA editing enzyme APOBEC1 and some of its homologs can act as DNA mutators. Mol Cell 10, 1247–1253.

6. Saraconi, G., Severi, F., Sala, C., Mattiuz, G. and Conticello, S.G. (2014) The RNA editing enzyme APOBEC1 induces somatic mutations and a compatible mutational signature is present in esophageal adenocarcinomas. Genome Biol 15, 417.

7. Niavarani, A., Shahrabi Farahani, A., Sharafkhah, M. and Rassoulzadegan, M. (2018) Pancancer Analysis Identifies Prognostic High-APOBEC1 Expression Level Implicated in Cancer In-Frame Insertions and Deletions. Carcinogenesis

8. Navaratnam, N., Morrison, J.R., Bhattacharya, S., Patel, D., Funahashi, T., Giannoni, F., Teng, B.B., Davidson, N.O. and Scott, J. (1993) The p27 catalytic subunit of the apolipoprotein B mRNA editing enzyme is a cytidine deaminase. J Biol Chem 268, 20709–20712.

9. Teng, B., Burant, C.F. and Davidson, N.O. (1993) Molecular cloning of an apolipoprotein B messenger RNA editing protein. Science 260, 1816–1819.

10. Blanc, V. and Davidson, N.O. (2010) APOBEC-1-mediated RNA editing. Wiley Interdiscip Rev Syst Biol Med 2, 594–602.

11. Skuse, G.R., Cappione, A.J., Sowden, M., Metheny, L.J. and Smith, H.C. (1996) The neurofibromatosis type I messenger RNA undergoes base-modification RNA editing. Nucleic Acids Res 24, 478–485.

12. Anant, S. and Davidson, N.O. (2000) An AU-rich sequence element (UUUN[A/U]U) downstream of the edited C in apolipoprotein B mRNA is a high-affinity binding site for Apobec-1: binding of Apobec-1 to this motif in the 3’ untranslated region of c-myc increases mRNA stability. Mol Cell Biol 20, 1982–1992.

13. Rosenberg, B.R., Hamilton, C.E., Mwangi, M.M., Dewell, S. and Papavasiliou, F.N. (2011) Transcriptome-wide sequencing reveals numerous APOBEC1 mRNA-editing targets in transcript 3’ UTRs. Nat Struct Mol Biol 18, 230–236.

14. Blanc, V., Park, E., Schaefer, S., Miller, M., Lin, Y., Kennedy, S., Billing, A.M., Ben Hamidane, H., Graumann, J., Mortazavi, A., Nadeau, J.H. and Davidson, N.O. (2014) Genome-wide identification and functional analysis of Apobec-1-mediated C-to-U RNA editing in mouse small intestine and liver. Genome Biol 15, R79.

15. Harjanto, D., Papamarkou, T., Oates, C.J., Rayon-Estrada, V., Papavasiliou, F.N. and Papavasiliou, A. (2016) RNA editing generates cellular subsets with diverse sequence within populations. Nat Commun 7, 12145.

16. Cole, D.C., Chung, Y., Gagnidze, K., Hajdarovic, K.H., Rayon-Estrada, V., Harjanto, D., Bigio, B., Gal-Toth, J., Milner, T.A., McEwen, B.S., Papavasiliou, F.N. and Bulloch, K. (2017) Loss of APOBEC1 RNA-editing function in microglia exacerbates age-related CNS pathophysiology. Proc Natl Acad Sci U S A 114, 13272–13277.

17. Rayon-Estrada, V., Harjanto, D., Hamilton, C.E., Berchiche, Y.A., Gantman, E.C., Sakmar, T.P., Bulloch, K., Gagnidze, K., Harroch, S., McEwen, B.S. and Papavasiliou, F.N. (2017) Epitranscriptomic profiling across cell types reveals associations between APOBEC1-mediated RNA editing, gene expression outcomes, and cellular function. Proc Natl Acad Sci U S A 114, 13296–13301.

18. Roth, S.H., Danan-Gotthold, M., Ben-Izhak, M., Rechavi, G., Cohen, C.J., Louzoun, Y. and Levanon, E.Y. (2018) Increased RNA Editing May Provide a Source for Autoantigens in Systemic Lupus Erythematosus. Cell Rep 23, 50–57.

19. Driscoll, D.M. and Casanova, E. (1990) Characterization of the apolipoprotein B mRNA editing activity in enterocyte extracts. J Biol Chem 265, 21401–21403.

20. Navaratnam, N., Shah, R., Patel, D., Fay, V. and Scott, J. (1993) Apolipoprotein B mRNA editing is associated with UV crosslinking of proteins to the editing site. Proc Natl Acad Sci U S A 90, 222–226.

21. Ikeda, T., Abd El Galil, K.H., Tokunaga, K., Maeda, K., Sata, T., Sakaguchi, N., Heidmann, T. and Koito, A. (2011) Intrinsic restriction activity by apolipoprotein B mRNA editing enzyme APOBEC1 against the mobility of autonomous retrotransposons. Nucleic Acids Res 39, 5538–5554.

22. Mehta, A., Kinter, M.T., Sherman, N.E. and Driscoll, D.M. (2000) Molecular cloning of apobec-1 complementation factor, a novel RNA-binding protein involved in the editing of apolipoprotein B mRNA. Mol Cell Biol 20, 1846–1854.

23. Lellek, H., Kirsten, R., Diehl, I., Apostel, F., Buck, F. and Greeve, J. (2000) Purification and molecular cloning of a novel essential component of the apolipoprotein B mRNA editing enzyme-complex. J Biol Chem 275, 19848–19856.

24. Lehmann, D.M., Galloway, C.A., Sowden, M.P. and Smith, H.C. (2006) Metabolic regulation of apoB mRNA editing is associated with phosphorylation of APOBEC-1 complementation factor. Nucleic Acids Res 34, 3299–3308.

25. Fossat, N., Tourle, K., Radziewic, T., Barratt, K., Liebhold, D., Studdert, J.B., Power, M., Jones, V., Loebel, D.A.F. and Tam, P.P.L. (2014) C to U RNA editing mediated by APOBEC1 requires RNA-binding protein RBM47. EMBO Rep 15, 903–910.

26. Yang, Y. and Smith, H.C. (1997) Multiple protein domains determine the cell type-specific nuclear distribution of the catalytic subunit required for apolipoprotein B mRNA editing. Proc Natl Acad Sci U S A 94, 13075–13080.

27. Blanc, V., Kennedy, S. and Davidson, N.O. (2003) A novel nuclear localization signal in the auxiliary domain of apobec-1 complementation factor regulates nucleocytoplasmic import and shuttling. J Biol Chem 278, 41198–41204.

28. Chester, A., Somasekaram, A., Tzimina, M., Jarmuz, A., Gisbourne, J., O’Keefe, R., Scott, J. and Navaratnam, N. (2003) The apolipoprotein B mRNA editing complex performs a multifunctional cycle and suppresses nonsense-mediated decay. EMBO J 22, 3971–3982.

29. Lau, P.P., Zhu, H.J., Baldini, A., Charnsangavej, C. and Chan, L. (1994) Dimeric structure of a human apolipoprotein B mRNA editing protein and cloning and chromosomal localization of its gene. Proc Natl Acad Sci U S A 91, 8522–8526.

30. Oka, K., Kobayashi, K., Sullivan, M., Martinez, J., Teng, B.B., Ishimura-Oka, K. and Chan, L. (1997) Tissue-specific inhibition of apolipoprotein B mRNA editing in the liver by adenovirus-mediated transfer of a dominant negative mutant APOBEC-1 leads to increased low density lipoprotein in mice. J Biol Chem 272, 1456–1460.

31. Navaratnam, N., Fujino, T., Bayliss, J., Jarmuz, A., How, A., Richardson, N., Somasekaram, A., Bhattacharya, S., Carter, C. and Scott, J. (1998) Escherichia coli cytidine deaminase provides a molecular model for ApoB RNA editing and a mechanism for RNA substrate recognition. J Mol Biol 275, 695–714.

32. Teng, B.B., Ochsner, S., Zhang, Q., Soman, K.V., Lau, P.P. and Chan, L. (1999) Mutational analysis of apolipoprotein B mRNA editing enzyme (APOBEC1). structure-function relationships of RNA editing and dimerization. J Lipid Res 40, 623–635.

33. Severi, F. and Conticello, S.G. (2015) Flow-cytometric visualization of C>U mRNA editing reveals the dynamics of the process in live cells. RNA Biol 12, 389–397.

34. Conticello, S.G., Ganesh, K., Xue, K., Lu, M., Rada, C. and Neuberger, M.S. (2008) Interaction between antibody-diversification enzyme AID and spliceosome-associated factor CTNNBL1. Mol Cell 31, 474–484.

35. Durocher, Y., Perret, S. and Kamen, A. (2002) High-level and high-throughput recombinant protein production by transient transfection of suspension-growing human 293-EBNA1 cells. Nucleic Acids Res 30, E9.

36. Klockenbusch, C. and Kast, J. (2010) Optimization of formaldehyde cross-linking for protein interaction analysis of non-tagged integrin beta1. J Biomed Biotechnol 2010, 927585.

37. Wolfe, A.D., Li, S., Goedderz, C. and Chen, X.S. (2020) The structure of APOBEC1 and insights into its RNA and DNA substrate selectivity NAR Cancer 2,

38. Webb, B. and Sali, A. (2016) Comparative Protein Structure Modeling Using MODELLER. Curr Protoc Bioinformatics 54, 5.6.1–5.6.37.

39. Wolfe, A.D., Arnold, D.B. and Chen, X. (2019) Comparison of RNA Editing Activity of APOBEC1-A1CF and APOBEC1-RBM47 Complexes Reconstituted in HEK293T Cells. J Mol Biol

40. Wong, L., Vizeacoumar, F.S., Vizeacoumar, F.J. and Chelico, L. (2021) APOBEC1 cytosine deaminase activity on single-stranded DNA is suppressed by replication protein A. Nucleic Acids Res 49, 322–339.

41. Brar, S.S., Sacho, E.J., Tessmer, I., Croteau, D.L., Erie, D.A. and Diaz, M. (2008) Activation-induced deaminase, AID, is catalytically active as a monomer on single-stranded DNA. DNA Repair (Amst) 7, 77–87.

42. Chelico, L., Sacho, E.J., Erie, D.A. and Goodman, M.F. (2008) A model for oligomeric regulation of APOBEC3G cytosine deaminase-dependent restriction of HIV. J Biol Chem 283, 13780–13791.

43. Chelico, L., Prochnow, C., Erie, D.A., Chen, X.S. and Goodman, M.F. (2010) Structural model for deoxycytidine deamination mechanisms of the HIV-1 inactivation enzyme APOBEC3G. J Biol Chem 285, 16195–16205.

44. McDougall, W.M., Okany, C. and Smith, H.C. (2011) Deaminase activity on single-stranded DNA (ssDNA) occurs in vitro when APOBEC3G cytidine deaminase forms homotetramers and higher-order complexes. J Biol Chem 286, 30655–30661.

45. Shlyakhtenko, L.S., Lushnikov, A.Y., Miyagi, A., Li, M., Harris, R.S. and Lyubchenko, Y.L. (2012) Nanoscale structure and dynamics of ABOBEC3G complexes with single-stranded DNA. Biochemistry 51, 6432–6440.

46. Shlyakhtenko, L.S., Lushnikov, A.Y., Miyagi, A., Li, M., Harris, R.S. and Lyubchenko, Y.L. (2013) Atomic Force Microscopy Studies of APOBEC3G Oligomerization and Dynamics. J Struct Biol 184, 217–225.

47. Bohn, M.-F., Shandilya, S.M.D., Silvas, T.V., Nalivaika, E.A., Kouno, T., Kelch, B.A., Ryder, S.P., Kurt-Yilmaz, N., Somasundaran, M. and Schiffer, C.A. (2015) The ssDNA Mutator APOBEC3A Is Regulated by Cooperative Dimerization. Structure 23, 903–911.

48. Feng, Y., Love, R.P., Ara, A., Baig, T.T., Adolph, M.B. and Chelico, L. (2015) Natural polymorphisms and oligomerization of human APOBEC3H contribute to single-stranded DNA scanning ability. J Biol Chem 290, 27188–27203.

49. Polevoda, B., McDougall, W.M., Tun, B.N., Cheung, M., Salter, J.D., Friedman, A.E. and Smith, H.C. (2015) RNA binding to APOBEC3G induces the disassembly of functional deaminase complexes by displacing single-stranded DNA substrates. Nucleic Acids Res 43, 9434–9445.

50. Pan, Y., Sun, Z., Maiti, A., Kanai, T., Matsuo, H., Li, M., Harris, R.S., Shlyakhtenko, L.S. and Lyubchenko, Y.L. (2016) Nanoscale Characterization of Interaction of APOBEC3G with RNA. Biochemistry 56, 1473–1481.

51. Wittkopp, C.J., Adolph, M.B., Wu, L.I., Chelico, L. and Emerman, M. (2016) A Single Nucleotide Polymorphism in Human APOBEC3C Enhances Restriction of Lentiviruses. PLoS Pathog 12, e1005865.

52. Adolph, M.B., Ara, A., Feng, Y., Wittkopp, C.J., Emerman, M., Fraser, J.S. and Chelico, L. (2017) Cytidine deaminase efficiency of the lentiviral viral restriction factor APOBEC3C correlates with dimerization. Nucleic Acids Res 45, 3378–3394.

53. Prochnow, C., Bransteitter, R., Klein, M.G., Goodman, M.F. and Chen, X.S. (2007) The APOBEC-2 crystal structure and functional implications for the deaminase AID. Nature 445, 447–451.

54. Shandilya, S.M.D., Nalam, M.N.L., Nalivaika, E.A., Gross, P.J., Valesano, J.C., Shindo, K., Li, M., Munson, M., Royer, W.E., Harjes, E., Kono, T., Matsuo, H., Harris, R.S., Somasundaran, M. and Schiffer, C.A. (2010) Crystal Structure of the APOBEC3G Catalytic Domain Reveals Potential Oligomerization Interfaces. Structure 18, 28–38.

55. Xiao, X., Li, S.-X., Yang, H. and Chen, X.S. (2016) Crystal structures of APOBEC3G N-domain alone and its complex with DNA. Nat Commun 7, 12193.

56. Bohn, J.A., Thummar, K., York, A., Raymond, A., Brown, W.C., Bieniasz, P.D., Hatziioannou, T. and Smith, J.L. (2017) APOBEC3H structure reveals an unusual mechanism of interaction with duplex RNA. Nat Commun 8, 1021.

57. Ito, F., Yang, H., Xiao, X., Li, S.-X., Wolfe, A., Zirkle, B., Arutiunian, V. and Chen, X.S. (2018) Understanding the Structure, Multimerization, Subcellular Localization and mC Selectivity of a Genomic Mutator and Anti-HIV Factor APOBEC3H. Sci Rep 8, 3763.

58. Matsuoka, T., Nagae, T., Ode, H., Awazu, H., Kurosawa, T., Hamano, A., Matsuoka, K., Hachiya, A., Imahashi, M., Yokomaku, Y., Watanabe, N. and Iwatani, Y. (2018) Structural basis of chimpanzee APOBEC3H dimerization stabilized by double-stranded RNA. Nucleic Acids Res

59. Shaban, N.M., Shi, K., Lauer, K.V., Carpenter, M.A., Richards, C.M., Salamango, D., Wang, J., Lopresti, M.W., Banerjee, S., Levin-Klein, R., Brown, W.L., Aihara, H. and Harris, R.S. (2017) The Antiviral and Cancer Genomic DNA Deaminase APOBEC3H Is Regulated by an RNA-Mediated Dimerization Mechanism. Mol Cell 69, 75–86.

60. Opi, S., Takeuchi, H., Kao, S., Khan, M.A., Miyagi, E., Goila-Gaur, R., Iwatani, Y., Levin, J.G. and Strebel, K. (2006) Monomeric APOBEC3G Is Catalytically Active and Has Antiviral Activity. J Virol 80, 4673–4682.

61. Opi, S., Kao, S., Goila-Gaur, R., Khan, M.A., Miyagi, E., Takeuchi, H. and Strebel, K. (2007) Human immunodeficiency virus type 1 Vif inhibits packaging and antiviral activity of a degradation-resistant APOBEC3G variant. J Virol 81, 8236–8246.

62. Shlyakhtenko, L.S., Lushnikov, A.J., Li, M., Harris, R.S. and Lyubchenko, Y.L. (2014) Interaction of APOBEC3A with DNA Assessed by Atomic Force Microscopy. PLoS One 9, e99354.

63. Lu, X., Zhang, T., Xu, Z., Liu, S., Zhao, B., Lan, W., Wang, C., Ding, J. and Cao, C. (2015) Crystal structure of DNA cytidine deaminase APOBEC3G catalytic deamination domain suggests a binding mode of full-length enzyme to single-stranded DNA. J Biol Chem 290, 4010–4021.

64. Chen, Z., Eggerman, T.L., Bocharov, A.V., Baranova, I.N., Vishnyakova, T.G., Csako, G. and Patterson, A.P. (2010) Hypermutation induced by APOBEC-1 overexpression can be eliminated. RNA 16, 1040–1052.

65. Petersen-Mahrt, S.K. and Neuberger, M.S. (2003) In vitro deamination of cytosine to uracil in single-stranded DNA by apolipoprotein B editing complex catalytic subunit 1 (APOBEC1). J Biol Chem 278, 19583–19586.

66. Morgan, H.D., Dean, W., Coker, H.A., Reik, W. and Petersen-Mahrt, S.K. (2004) Activation-induced cytidine deaminase deaminates 5-methylcytosine in DNA and is expressed in pluripotent tissues: implications for epigenetic reprogramming. J Biol Chem 279, 52353–52360.

67. Ikeda, T., Ohsugi, T., Kimura, T., Matsushita, S., Maeda, Y., Harada, S. and Koito, A. (2008) The antiretroviral potency of APOBEC1 deaminase from small animal species. Nucleic Acids Res 36, 6859–6871.

68. Severi, F., Chicca, A. and Conticello, S.G. (2011) Analysis of Reptilian APOBEC1 Suggests that RNA Editing May Not Be Its Ancestral Function. Mol Biol Evol 28, 1125–1129.

69. Caval, V., Jiao, W., Berry, N., Khalfi, P., Pitré, E., Thiers, V., Vartanian, J.-P., Wain-Hobson, S. and Suspène, R. (2019) Mouse APOBEC1 cytidine deaminase can induce somatic mutations in chromosomal DNA. BMC Genomics 20, 858.

70. Feldser, D.M., Hackett, J.A. and Greider, C.W. (2003) Telomere dysfunction and the initiation of genome instability. Nat Rev Cancer 3, 623–627.

71. Aoufouchi, S., Faili, A., Zober, C., D’Orlando, O., Weller, S., Weill, J.-C. and Reynaud, C.-A. (2008) Proteasomal degradation restricts the nuclear lifespan of AID. J Exp Med 205, 1357–1368.

72. Mondal, S., Begum, N.A., Hu, W. and Honjo, T. (2016) Functional requirements of AID’s higher order structures and their interaction with RNA-binding proteins. Proc Natl Acad Sci U S A 113, E1545–54.

