## Supplementary Figures for "Dimerisation of APOBEC1 is dispensable for its RNA/DNA editing activity and modulates its availability"

**Supplementary Figure 1**

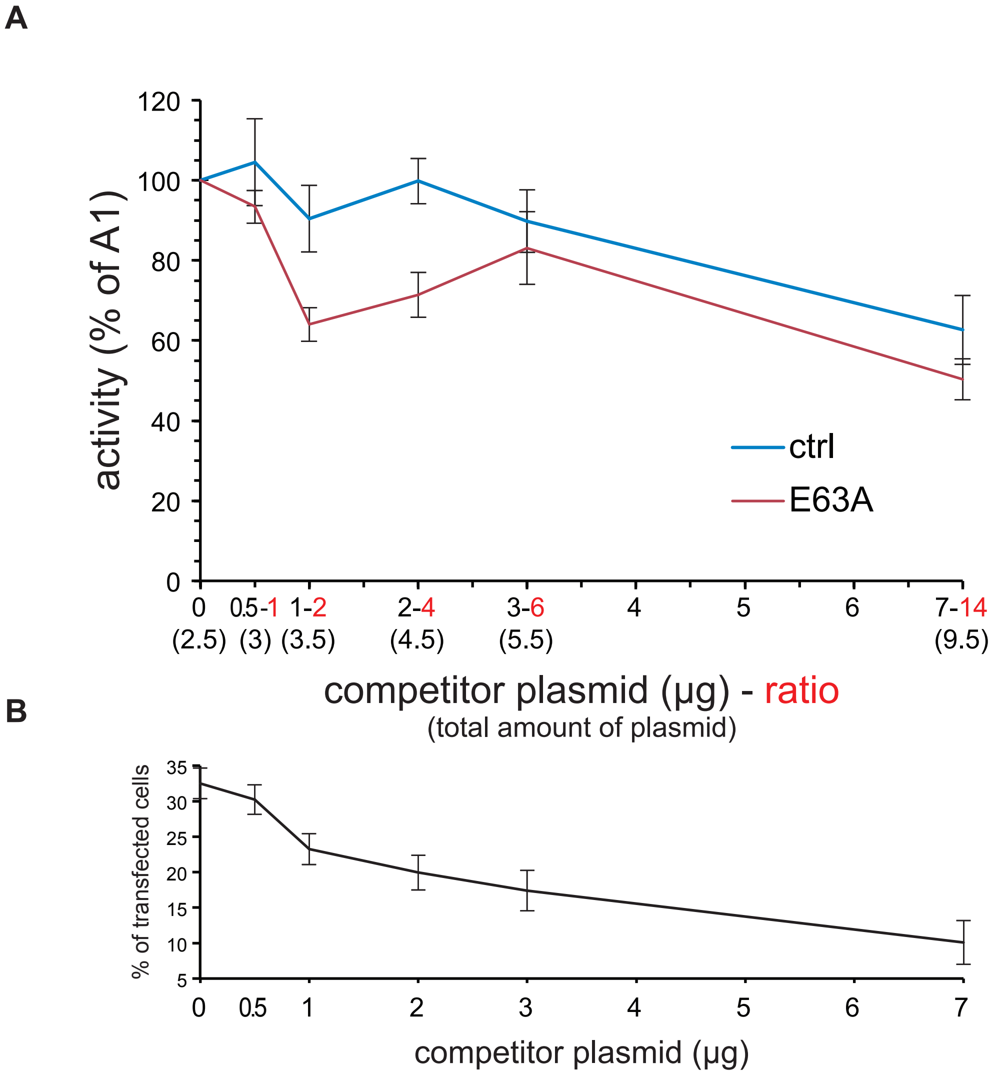


**Linearity of APOBEC1 activity with increasing amount of plasmid DNA**

RNA editing of rat APOBEC1 in HEK293T cells transiently cotransfected with plasmids encoding for the mCherry-ApoB-EGFP reporter (1 µg), APOBEC1 (0.5 µg), A1CF (1 µg), and increasing amounts of either a control plasmid (ctrl) or a catalytically inactive APOBEC1 mutant (E63A). (A) the panel shows the APOBEC1-dependent RNA editing of the reporter (percentage of gated cells) normalised to the activity of APOBEC1 in absence of competitor DNA. The ratio of the competitor DNA with APOBEC1 is shown in red, and the amount of total DNA used in the transfection is shown in parentheses. (B) The lower panel shows the transfection efficiency (mCherry(+) cells) for each point.The error bars represent the standard deviation from three experiments.

### Supplementary Figure 2

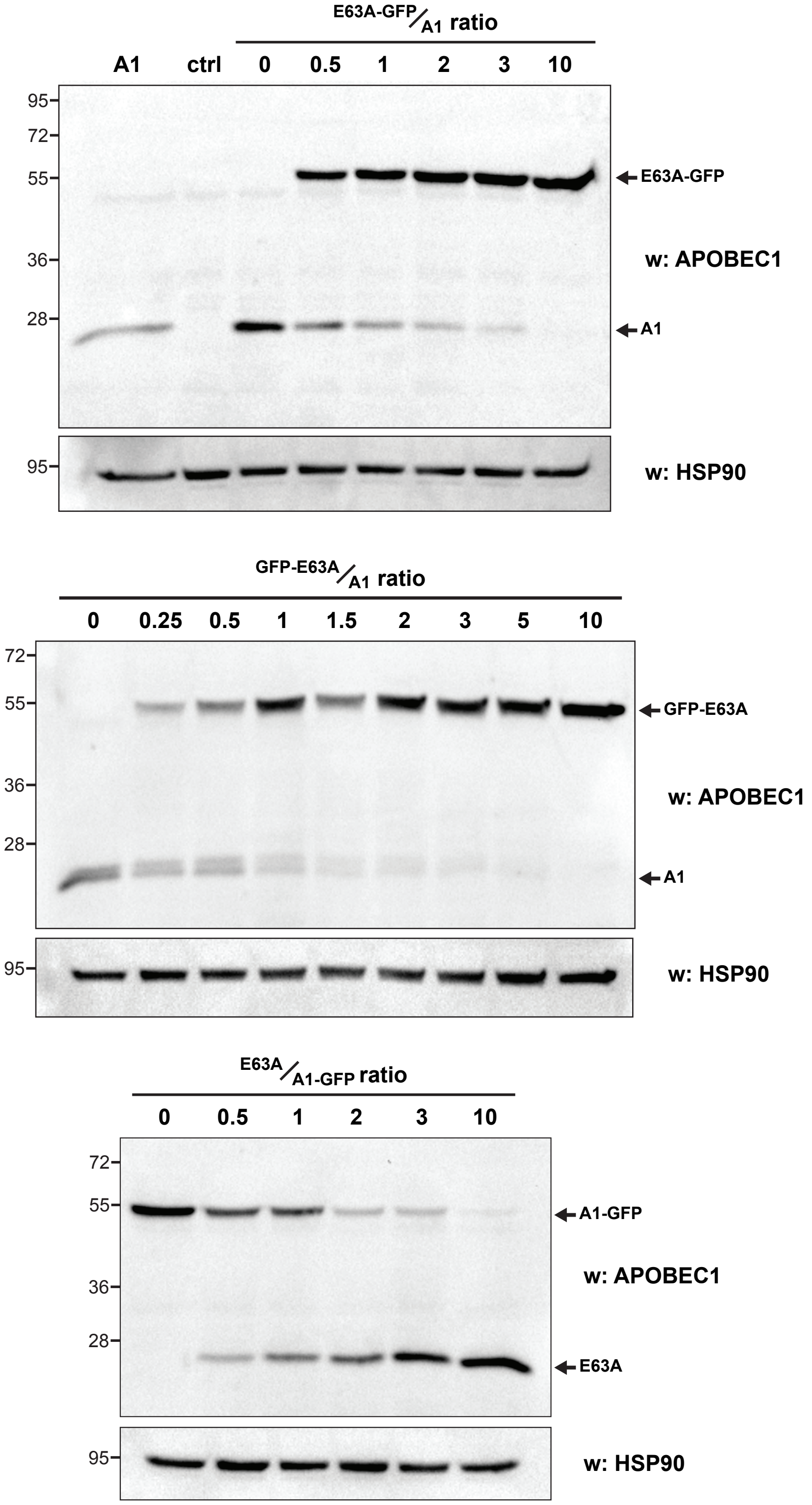


**Coexpression of different ratios of APOBEC1 constructs**

Representative western blots showing the expression levels various combinations of APOBEC1 constructs in transiently transfected HEK293T cells. 2 µg of total plasmid DNA were transfected in cells using the ratios indicated on the blots. There is not representative blot for GFP-A1/A1-GFP cotransfections, as the identical molecular weight does not allow their discrimination.

### Supp.Fig 3

**A**

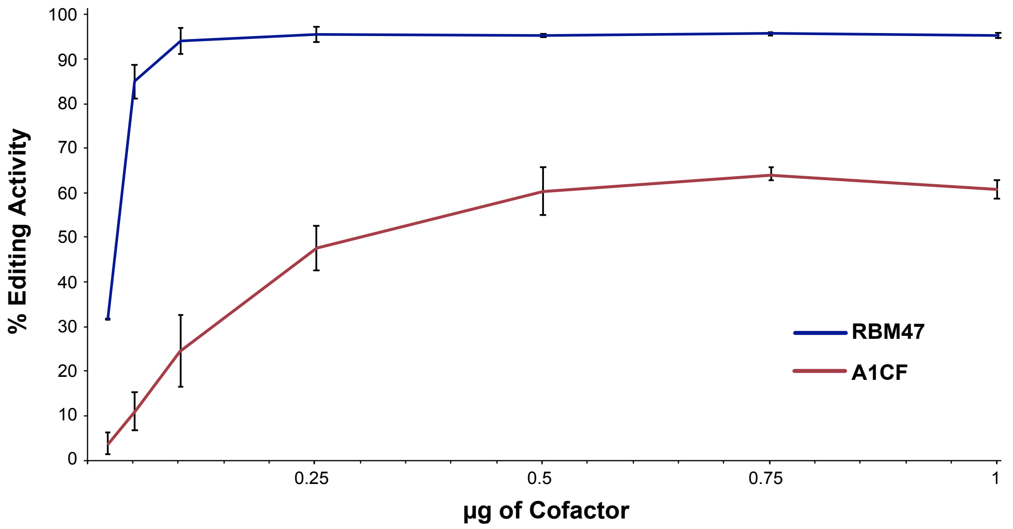


**B**

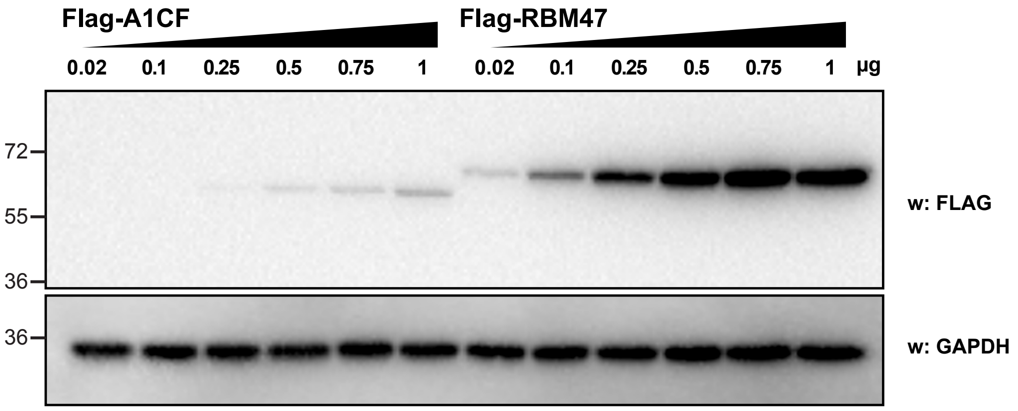


**Supplementary Figure 3. (A) RNA Editing Activity of APOBEC1 in presence of either RBM47 or A1CF and RBM47 cofactors.** Increasing amounts (20 ng - 1 µg) of A1CF (blue) and RBM47 (red) were cotransfected in HEK293T cells together with 1 µg of editing reporter plasmid and 1 µg of APOBEC1. (B) Comparison of the expression levels of the cofactors. Increasing amounts (20 ng - 1 µg) of FLAG-tagged A1CF and RBM47 were cotransfected in HEK293T cells and their expression was assayed by western blot using anti-FLAG antibody.

### Supplementary Figure 4

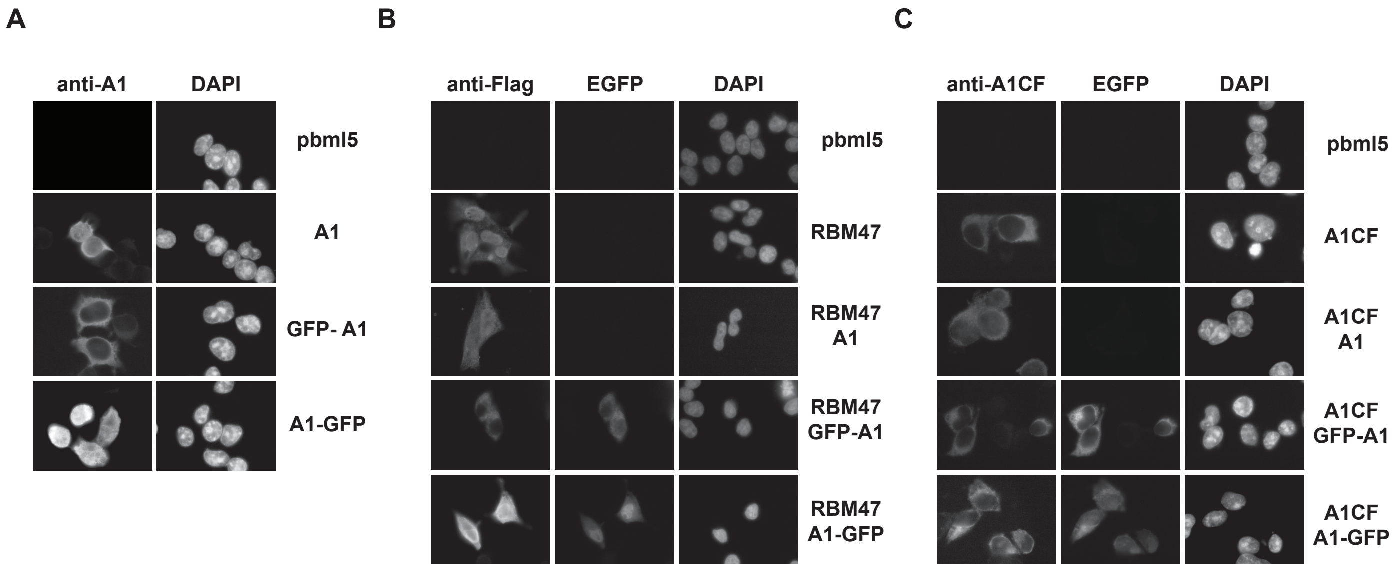


**Supplementary Figure 4. Localisation of APOBEC1 (A), and of RBM47 (B) or A1CF (C) in presence of the APOBEC1 constructs.** Representative images of HEK293T cells transfected with either RBM47 or A1CF alone or in combination with the APOBEC1 constructs (A1, GFP-A1, A1-GFP). An empty plasmid was used as a negative control. Localisation of the various proteins was assessed by GFP fluorescence for the GFP-tagged APOBEC1 chimeras, and by anti-FLAG or anti-A1CF antibodies for visualization of APOBEC1, RBM47 or A1CF, respectively.

### Supplementary Figure 5

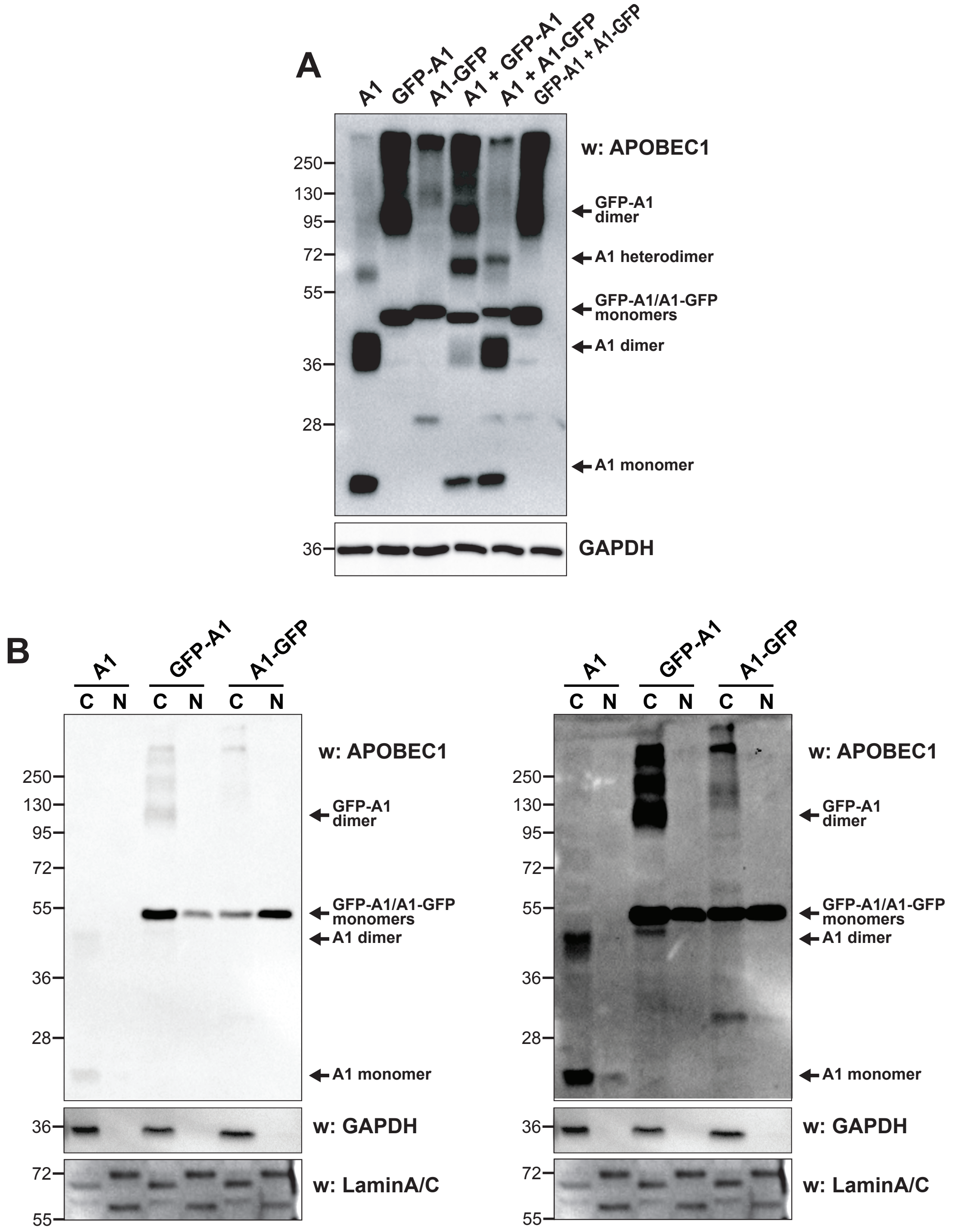


**Supplementary figure 5: Different exposures of the crosslinking experiments shown in Figure 3B (A) and 3C (B).**

### Supplementary Figure 6

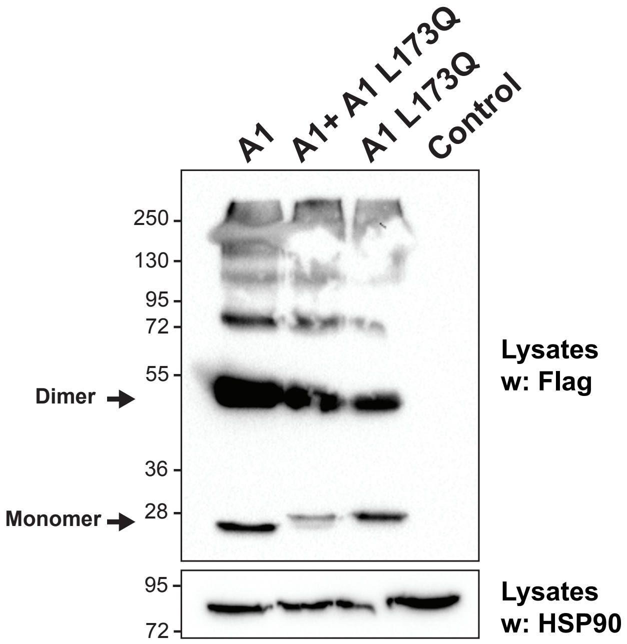


**Supplementary Figure 6. Crosslinking of APOBEC1 L173Q in whole cell lysates.** HEK293T cells were transiently cotransfected as indicated and were treated with 2% formaldehyde (10' at room temperature). The crosslinking was blocked with 1.25M glycine and cells were then lysed. Following SDS/PAGE, western blots were probed with anti-APOBEC1 antibodies.

### Supplementary Figure 7

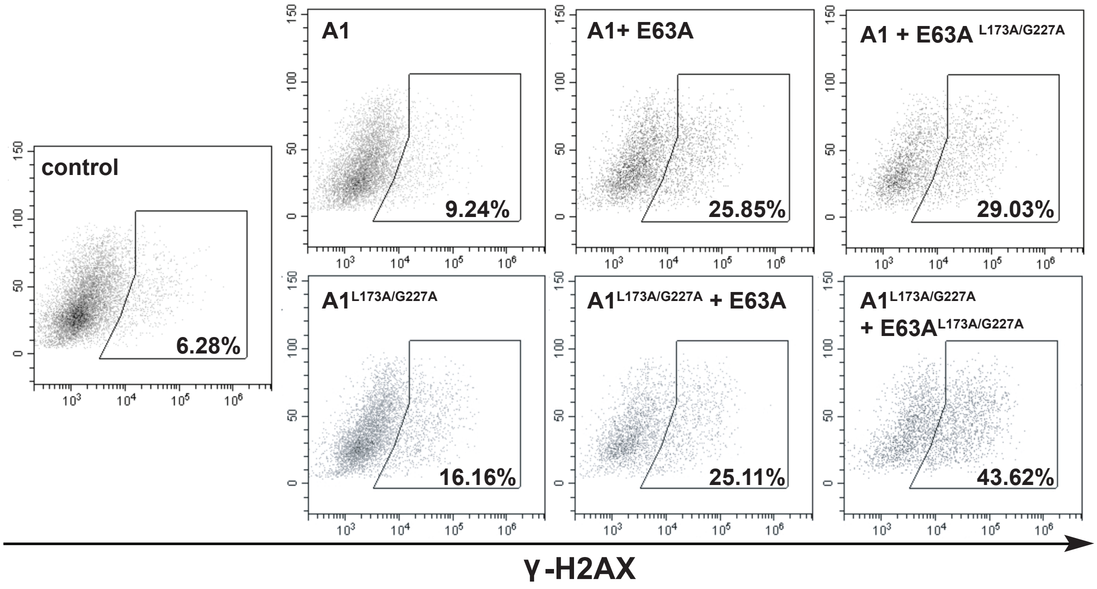


**Supplementary Figure 7. Representative FACS profiles of transfected cells after intracellular staining with anti-γ-H2AX antibody.** Induction of DNA damage in HEK-293T cells transfected with either APOBEC1 (A1) or APOBEC1$^{L173A/G227A}$ (L173A/G227A) and an excess of catalytically inactive APOBEC1 (E63A or E63/L173A/G227A). Cells were transfected with a plasmid mix containing catalytically active and inactive APOBEC1 constructs in a 1:3 ratio (2 µg). In all experiments shown a cotransfected EGFP expressing plasmid (100ng) was used to identify transfected cells.
